# The genetic basis of adaptation in phenology in an introduced population of Black Cottonwood (*Populus trichocarpa*, Torr. & Gray)

**DOI:** 10.1101/2020.06.17.156281

**Authors:** Rami-Petteri Apuli, Thomas Richards, Martha Rendon, Almir Karacic, Ann-Christin Rönnberg Wästljung, Pär K. Ingvarsson

## Abstract

- Entering and exiting winter dormancy presents important trade-offs between growth and survival at northern latitudes and many forest trees display local adaptation across latitude. Transfers of a species outside its native range introduce the species to novel combinations of environmental conditions potentially requiring different combinations of alleles to optimize growth.
- We performed genome wide association analyses and a selection scan in a *P. trichocarpa* mapping population derived from crossings between clones collected across the native range and introduced into Sweden. GWAS analyses were performed using phenotypic data collected across two field seasons and in a controlled phytotron experiment.
- We uncovered 629 putative candidate genes associated with spring and autumn phenology traits as well as with growth. Many regions harboring variation significantly associated with the initiation of leaf shed and leaf autumn coloring appeared to have been evolving under positive selection in the native environments of *P. trichocarpa*.
- A comparison between the candidate genes identified with results from earlier GWAS analyses performed in the native environment found a smaller overlap for spring phenology traits than for autumn phenology traits, aligning well with earlier observations that spring phenology transitions have a more complex genetic basis that autumn phenology transitions.

## Introduction

At northern latitudes winter conditions are unfavorable to active plant growth through a combination of low temperatures, frosts, and light conditions leading perennial plants to avoid these conditions by entering winter dormancy (hereafter dormancy) (Doorenbos, 1953). Transitions from active growth to dormancy and from dormancy to active growth are controlled by different environmental cues where the transition to dormancy is primarily induced by changes in photoperiod (Fracheboud *et al.*, 2009) or light quality (Clapham *et al.*, 1998), while the release of dormancy is induced by prolonged exposure to low temperatures followed by increasing temperatures reactivating growth (Singh *et al.*, 2018). Incorrect timing of phenology transitions is known to result in loss of potential growth through extended dormancy or loss of realized growth in the form of damage to important tissues such as meristems and leaves from exposure to unfavorable conditions. Dormancy hence represents an important life history trade-off between growth and survival and maladapted individuals are therefore likely to suffer lowered reproductive success and/or biomass production, both of which may have large ecological and economic repercussions (Loehle, 1998).

Populations of widespread species often display signatures of phenotypic and genetic adaptation to their native environments, even in species with considerable gene flow among the populations (Howe *et al.*, 2003). This phenomenon, known as local adaptation, often arises from positive selection (Fournier-Level *et al.*, 2011), which leaves distinct and detectable signatures across the genome (Maynar Smith & Haigh, 1974; Begun & Aquadro, 1992). The strength of selection along with local rates of recombination and gene flow are the major determining factors of the extent and magnitude of signatures of selection (Begun & Aquadro, 1992; Vitti *et al.*, 2013). Furthermore, associations between segregating polymorphisms in the genomes of individuals and their phenotypes or measures of their natal environment can be explored by performing genome wide association studies (GWAS) (Josephs *et al.*, 2017). Local adaptation to climate and photoperiod has been observed in a large number of species with wide South-North distribution ranges including *Arabidopsis thaliana* (Fournier-Level *et al.*, 2011), *Picea abies* (Beuker, 1994) and *Populus tremula* (Luquez *et al.*, 2008). Empirical studies suggest that the genetic basis of local adaptation can be highly polygenic, where a majority of the loci and alleles conferring local adaptation have small effects (Rockman, 2012) although large effect loci have been observed in some systems (e.g. Wang *et al.*, 2018). Due to the polygenic nature of these traits, the genetic architecture of local adaptation to climate can be very diverse among even closely related species, despite the adaptation being driven by very similar environmental conditions (e.g McKown *et al.*, 2014; Wang *et al.*, 2018)

Species within the genus *Populus* are deciduous, early succession trees with wide distributions across the northern hemisphere, spanning from the equator to the northern limits of tree growth. The rapid growth rate and ability to generate natural clones (Taylor, 2002; Lin *et al.*, 2018) has spurred economic interest in the genus (Taylor, 2002), while many of the species in the genus are also considered keystone species in their natural habitats (Kouki *et al.*, 2004). *Populus* species are frequently utilized in biomass production for forest industry, even outside of their natural distribution ranges (Dickmann & Kuzovkina, 2014). In northern Europe, biomass production with *Populus* species is an underutilized option due to the phenological maladaptation of commercially bred varieties (Karacic *et al.*, 2003). Commercial interest thus exists for adapting non-native *Populus* species to growth under northern European conditions. Black cottonwood (*Populus trichocarpa*) is a deciduous tree native to North America with continuous distribution in western and northwest North America from California to Alaska. The species has been thoroughly studied in its natural range and has been found to display signatures of local adaptation to climate and photoperiod across its natural range (Evans *et al.*, 2014; McKown *et al.*, 2014). However, exploring the genetic architecture of these traits under novel conditions could reveal novel genes associated with them and comparisons with the results from the natural range present an opportunity to uncover further details of adaptation in these traits.

In this study, we perform a field trial and a controlled environment phytotron trial to collect data on spring and autumn phenology and growth traits in *P. trichocarpa*. We use the data to dissect the genetic basis of these traits using both population genomic approaches and genome-wide association studies (GWAS). We compare the candidate genes identified across two successive years to either identify genes that appear to have reliable effects on different traits or to identify genes with possible pleiotropic effects across multiple traits. We also compare candidate genes identified under field conditions and the phytotron experiment to explore the genetic relationships between different measures of bud burst and between the bud set proxy traits, leaf senescence and autumn coloring, and bud set. Finally, we compare the identified candidate genes with results from earlier studies performed in the natural range of *P. trichocarpa* to explore the nuances of genetic control of the phenology traits under native and novel environments

## Materials and Methods

### Plant material, phenotyping of field experiment and climate data

The trees used in this study are first- or second-generation offspring generated from crosses between *P. trichocarpa* trees collected from across the natural range in western North America (Table S1). Individuals with high growth and well-adapted phenology timing were then chosen from 34 families with 1 to 21 full-sibs per family. Chosen individuals were then clonally replicated and planted in 2003 in five complete blocks together with some of the parent trees at Krusenberg near Uppsala, Sweden (59°44’44.2“N 17°40’31.5“E). At the time of this study (2017-2018), 564 live ramets from 120 unique genotypes remained at the field site, all of which were genotyped. Climate data for the field site was obtained from the SLU Ultuna climate station (Table S2).

The trees were phenotyped every 2-5 days in two successive years, 2017 (17) and 2018 (18), for bud burst (BB) (spring phenology) and for leaf shed (LS) and autumn coloring (CO) (autumn phenology). Leaf shed and autumn coloring were measured as proxies for bud set (BS), which is the most relevant measurement of season-ending growth (Fracheboud *et al.* 2009), but which is hard to measure accurately in fully grown trees. The diameter at breast height (DBH) of the trees was measured in 2017 and was used as a proxy for lifetime growth.

Bud burst was scored using a scale with six steps, ranging from fully dormant buds (1) to fully opened buds with unfurled leaves and active shoot growth (6) (Table S3). Leaf shed was scored on a scale ranging from 1 to 5, with each stage describing a window of 20% of leaves shed (stage 1: 0% to 20% leaves shed, stage 5: 80% to 100% leaves shed). Autumn coloring was measured slightly differently between the two years, using a scale ranging from 1 to 5 in 2017 and a scale ranging from 1 to 8 in 2018 (Table S4), based on the level of yellowing in the leaf crown with 1 being fully green leaves and 5/8 being fully yellow leaves.

### Plant material, conditions and phenotyping of phytotron experiment

In February of 2018, cuttings were taken from 99 clones in the Krusenberg field trial. These cuttings were taken from stems of root suckers growing from stumps of thinned individuals or from branches of mature trees and stored at −4 °C until planted in pots in March of 2018. Two cuttings of similar length (~10 cm) from the same clone were planted in each pot. This was done in three replicates to produce a 3-block randomized design. After sprouting, the less vigorous cutting was removed leaving only one cutting per pot. Cuttings were then put through two simulated seasons described in Table S5.

Bud burst and bud set were measured during the second simulated season (Table S5). Bud burst was scored following the six-step scale used in the field trial (Table S3), but the buds on the saplings were divided into four classes and scored separately. These classes were the apical bud on the longest stem (top), the branch buds (brn) consisting of all the buds on lateral branches if present, the top 50 % of the stem buds (stt) and the bottom 50 % of the stem buds (stb) consisting of all the buds on the top and the bottom half of the main stem respectively. Bud set was scored in the highest situated undamaged bud (apical if possible, see below) following a seven-stage scale introduced in Rohde *et al.*, (2011) with minor changes in stage numbering, ranging from growing apical meristem (1 in our numbering, 3 in Rohde *et al.*, 2011) to fully set bud (7 in our numbering, 0 in Rohde *et al.*, 2011).

### Genotyping, SNP calling and filtering

Leaf samples were collected from trees in the Krusenberg trial in the autumn of 2016 and stored dried with silica gel until DNA extractions. DNA extractions were repeated for a few clones using leaves taken from the cuttings used for the phytotron experiment. Genomic DNA was extracted using a Macherey-Nagel NucleoSpin® Plant II kit according to manufacturer’s instructions. Quality and concentrations of the DNA was assessed using a NanoDrop spectrophotometer. Paired-end sequencing libraries with insert sizes of 350bp were constructed for all samples at the National Genomics Infrastructure at the Science for Life Laboratory in Stockholm, Sweden. Whole-genome sequencing with a target depth of 20× was performed using an Illumina HiSeq X platform with 2×150-bp paired-end reads.

Sequencing reads for all accessions were mapped against the reference genome of *P. trichocarpa* v3.0, using BWA-MEM (v0.7.17) (Li & Durbin, 2009) using default parameters. Depth and breadth of coverage were assessed in order to confirm that all samples had a minimum coverage of 10X (range from 12 to 49X, see Table S6) Post-mapping filtering removed unmapped reads (samtools v1.10) (Li *et al.*, 2009) and tagged duplicate reads (picard MarkDuplicates v2.10.3) (http://broadinstitute.github.io/picard/), which did not exceed 14% of the libraries (ranging from 3 to 13.8%).

We used GATK v3.8 (Van der Auwera *et al.*, 2013) to call variants. We performed local realignment around indels with RealignerTargetCreator and IndelRealigner (default parameters). Sample variants were called using HaplotypeCaller, producing gVCF files (−ERC GVCF). Samples were hierarchically merged into intermediate gVCF files using CombineGVCFs and were finally called jointly with GenotypeGVCFs. SNPs were selected using SelectVariants and filtered with VariantFiltration (QD < 2.0; FS > 60.0; MQ < 40.0; ReadPosRankSum < −8.0; SOR > 3.0; MQRankSum < −12.5). SNPs were pruning with vcf/bcftools (Li *et al.*, 2009; Danecek *et al.*, 2011) to remove positions with extreme depth (min-meanDP 16, max-meanDP 33; these thresholds correspond to the average depth ± one standard deviation), missing in more that 30% of the samples, non-biallelic SNPs with minor allele frequencies < 0.05, or SNPs displaying an excess of heterozygosity (FDR <0.01). This resulted in a data set consisting of 7,297,862 SNPs. SNPs were further filtered for allele number = 2, minor allele frequency (MAF) > 0.05 and extreme deviation from Hardy-Weinberg equilibrium (HWE) < 10^−6^. After filtering, 7,076,549 SNPs were retained and used in all downstream analyses.

### Missing data and phenotypic data imputation

During the field experiment individual trees sometimes passed through more than one phenology stage between two successive phenotypings. To account for these missing phenotypes, we first converted each ordinal stage into the number of Julian days (measured from January 1) when a stage was first observed for a given individual tree. This was done to remove accidental reversals of phenology stages which occurred at low frequencies in the data set due differences in subjective scoring by different observers or through shedding of yellowed leaves which sometimes case an apparent ‘greening’ of some trees. Once an individual tree had transitioned to the next phenology stage the Julian date was recorded for that stage and any subsequent reversals were discarded. For the first (1) stage for both spring and autumn phenology we used the last observed date as the observation point, as the first stage denotes ‘no change’ making earlier observations of the stage uninformative about the progress of phenology.

A local regression model (LOESS) was fitted through the transition days to estimate the missing stage transition days for each individual separately. The method fits a non-linear curve that is not constrained to fit any a-priori distribution for each individual separately. This allows estimation of the day in which these individuals entered each ordinal developmental stage allowing us to include individuals not observed at transitions between stages in the downstream analyses (For more information see Richards *et al.*, 2020)). The mean of each clone was then calculated and used as the stage-specific phenotype for the genotype in all subsequent analyses. We observed negligible differences between estimated BLUPs and means (Fig. S1) due to the simplicity of our experimental design.

### Choice of stages

The phenology traits were phenotyped using multiple stages that are highly correlated across individuals (Fig. S2). Most information is conferred by the initial transition (stage 1 to 2) and final transition stages in terms of growth period and vulnerability to damage and we therefore chose these two stages to serve as representative time points for phenology transitions in our data. For bud set and leaf shed, the stages used were the second and the last stage. For autumn coloring, the last stage was chosen to represent the end of the phenology transition, but here we instead used the third stage to represent the initiation stage, as the third stage was directly phenotyped in both years (Table S4). For bud burst, the start of the second stage was chosen as the beginning of phenology transition, and the fourth stage was chosen as the end of transition as this represents the stage when leaves begin to emerge and unfurl beginning the active photosynthesis.

For the field phenology traits, a Welch two sample t-test was performed to confirm whether the differences observed in phenology timing between the two years was statistically significant. The narrow sense heritability (h^2^) was also calculated for each of our chosen traits using the “heritability” package (Kruijer *et al.*, 2014) in R (R Core Team, 2014), a marker-based method developed specifically for plant data, utilizing the standardized kinship matrix calculated in GEMMA (Zhou & Stephens, 2012) and using the imputed phenotypes from each individual as replicates for each clone to estimate to what extent phenotypic variation was heritable (Table S7) (Kruijer *et al.*, 2014).

### GWAS and Lindley score

A genome wide association study was performed for each of the 23 chosen traits utilizing a univariate linear mixed model implemented in with GEMMA (v. 0.98.1). All field traits were run with no covariates, but for traits measured in the phytotron, a binary covariate was included to indicate the status of the apical meristem (damaged/not damaged) to account for effects of the phytotron issues (see Results). To take advantage of the large number of markers available and better utilize the information contained in the linkage disequilibrium among adjacent markers, we used the Lindley score-based method introduced in Bonhomme *et al.*, (2019). The local Lindley score is calculated using information derived from multiple adjacent SNPs, thereby limiting the number of tests performed while utilizing all the available data. Each *p*-value that exceeds a user set threshold (ξ, on a logarithmic scale) will contribute positively to a local Lindley score and vice versa for SNPs that fall below the threshold. Lindley scores can not to go below zero regardless of how many p-values that fall below a given threshold. If enough adjacent tests are significant or a single test is highly significant, the local Lindley score will rise above the chromosome specific significance level2, signifying an area of interest, which is especially useful for highlighting areas containing multiple weakly significant markers. The Lindley score is the result of a directional process and the leading-edge slope (hereafter slope) is the area of interest.

### LD decay, candidate genes and candidate gene comparisons

To identify candidate genes around the significant slopes revealed by the local Lindley score analyses, the rate of decay of linkage disequilibrium (LD) was estimated following Wang *et al.*, (2016). Briefly, SNP markers were randomly thinned down to 100,000 markers using PLINK 1.9 (Purcell *et al.*, 2007) and remaining markers were used to calculate the squared correlation coefficients (r^2^) between all SNP pairs in non-overlapping 50 kbp windows using PLINK 1.9. The decay of LD across physical distance was then estimated using nonlinear regression of pairwise *r*^2^ against physical distance between sites in base pairs (Remington *et al.*, 2001). LD decays over, on average, 10 kbp (Fig. S3) in our *P. trichocarpa* population, and we used this information to determine putative candidate genes for the regions that showed significant association in the GWAS. Boundaries of significant slopes were extended by ±10 kbp to search for genes using the *P. trichocarpa* v3.1 annotation (Tuskan *et al.*, 2006) available at Phytozome 12 (https://phytozome.jgi.doe.gov/pz/portal.html).

The sets of candidate genes for each trait were compiled and yearly comparisons between traits and between the different bud set traits from the phytotron experiment were illustrated using Venn diagrams. We compared candidate genes identified for spring and autumn phenology with candidate genes identified in two earlier studies in *P. trichocarpa*, Evans *et al.*, (2014) and McKown *et al.*, (2014), to reveal any similarities in candidate genes identified based on common garden data from Sweden, Canada or the phytotron experiment.

GO-term enrichment analysis was performed on each of the study traits by merging the candidate genes from each stage of the trait. The analysis was performed online at PopGenIE (https://popgenie.org) and was performed on the *Arabidopsis thaliana* synonyms of the genes using the default settings of the tool. PopGenIE uses Fisher’s exact test with False Discovery Rate (FDR) correction with a corrected p-value threshold of 0.05 and minimum of two genes by default.

### Signatures of positive selection

We calculated two haplotype-based test statistics to detect positive selection, the integrated haplotype score iHS, (Voight *et al.*, 2006) and H12, which measures haplotype homozygosity and is especially useful for finding soft selective sweeps (Garud *et al.*, 2015). Both test statistics were calculated using selscan v1.2.0a (Szpiech & Hernandez, 2014). The genetic map positions of all SNP markers were calculated based on the population averaged recombination rates estimated using LDhat (Wang *et al.*, 2016) with missing values set to zero. iHS does not produce estimates for zero values leading to slightly different numbers of estimates between the two methods. For both statistics the top 0.1 percentile was used as a threshold for signatures of selection. To test for possible enrichments between SNPs showing evidence for positive selection and significant SNPs from the GWAS, we used hypergeometric distribution tests on each trait separately. Peak SNP’s were then pinpointed for each selection scan using the ggpmisc package (Aphalo 2020) in R. A peak was the SNP with highest selection scan score of all SNP’s within a window of 20001 bp centered at that SNP. Only the peaks in the top 0.1 percentile (hereafter top peaks) were used for downstream analyses for both estimates. Genes within 10 kbp of the top peaks were then identified and compared with our GWAS candidate genes. Genes surrounding the selection peaks were analyzed for GO-enrichments using the default settings on PopGenIE enrichment analysis tool.

## Results

### Phenotypic variation and heritability

All traits display variation both within and between years, though BB2-top (Fig. 1e) and BS2 (Fig. 1f) are noticeably less variable than the other traits (Fig. 1). In the field traits there are highly significant (p < 0.001) differences between years 2017 and 2018. All chosen traits with the exception of BB2-top and BS2 had an appreciable level of heritability (h^2^ > 0.15) with only BB2-top having a heritability less than 0.05 (Table S7). None of the traits could be considered to be identical, though BB2-stt and BB4-stt, BB4-stt and BB4-stb, and CO3-17 and CO8-17 all had high correlations (r^2^ > 0.8) (Fig. S2).

**Figure 1:**
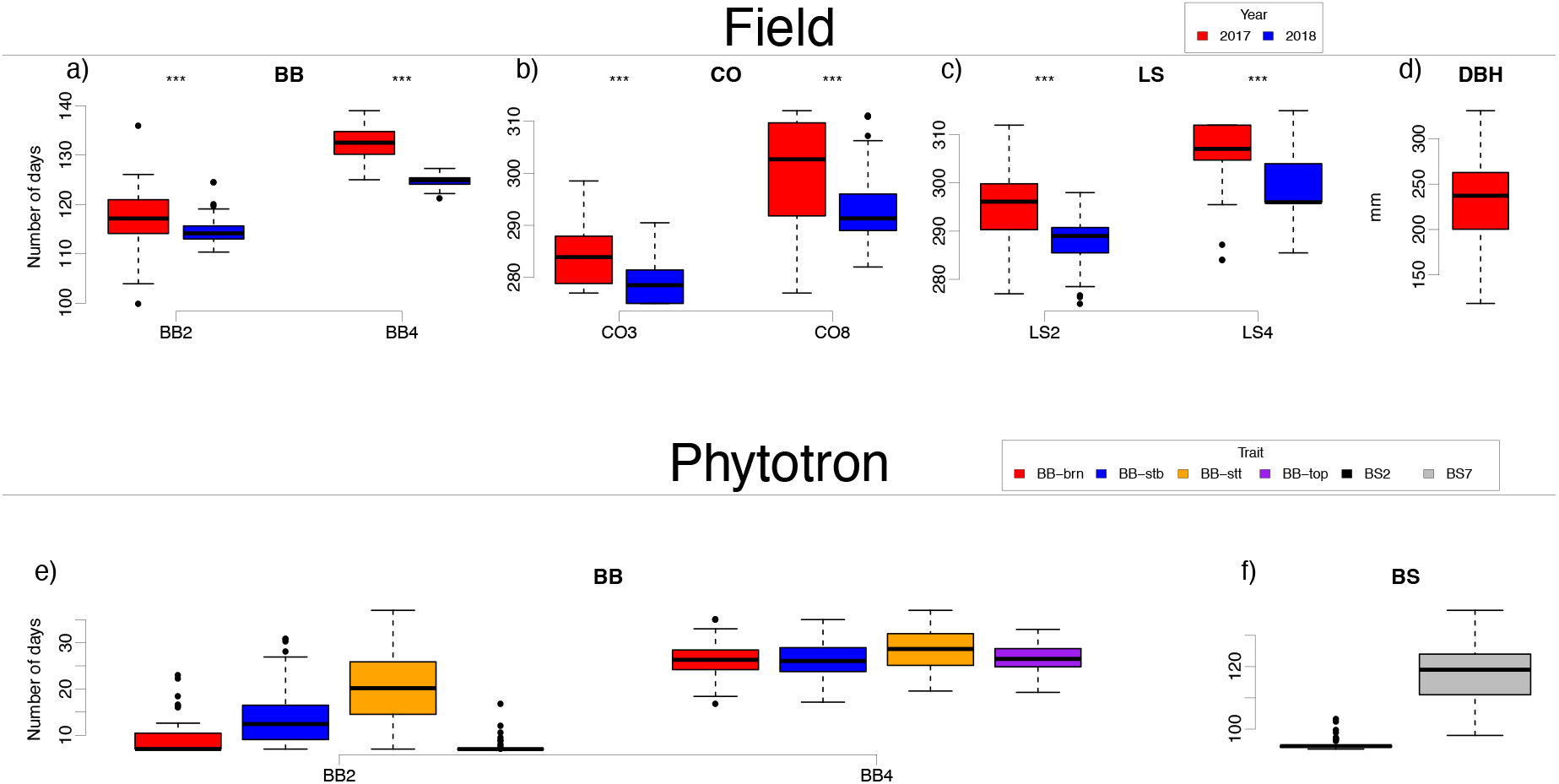
Phenotypes of a) bud burst (BB), b) autumn coloring (CO), c) leaf shed (LS), d) diameter at breast height (DBH) in the field and e) bud burst and f) bud set (BS) in phytotron.

The initiation of bud burst was visibly different between the different parts of the plant in the phytotron experiment (brn, stb, stt and top). However, these differences largely disappeared by the 3^rd^ stage for brn, stb and top, while stt remained different until the 4^th^ stage. At stage 5, the different parts of the plants were all behaving in a similar fashion (Fig. 2a). In July/August of 2018, the phytotron equipment malfunctioned causing the simulated winter temperatures to shift into unexpectedly high temperatures, causing early bud flush and subsequent bud damage in some of the cuttings once the winter conditions were restored. When individuals with a damaged or absent apical bud were compared, bud burst timing of both stt (Fig. 2b) and stb (Fig. 2c) were significantly different whereas branches showed no difference (Fig. 2d). A significant difference was also seen in the latter stages of bud set (Fig. 2e).

**Figure 2:**
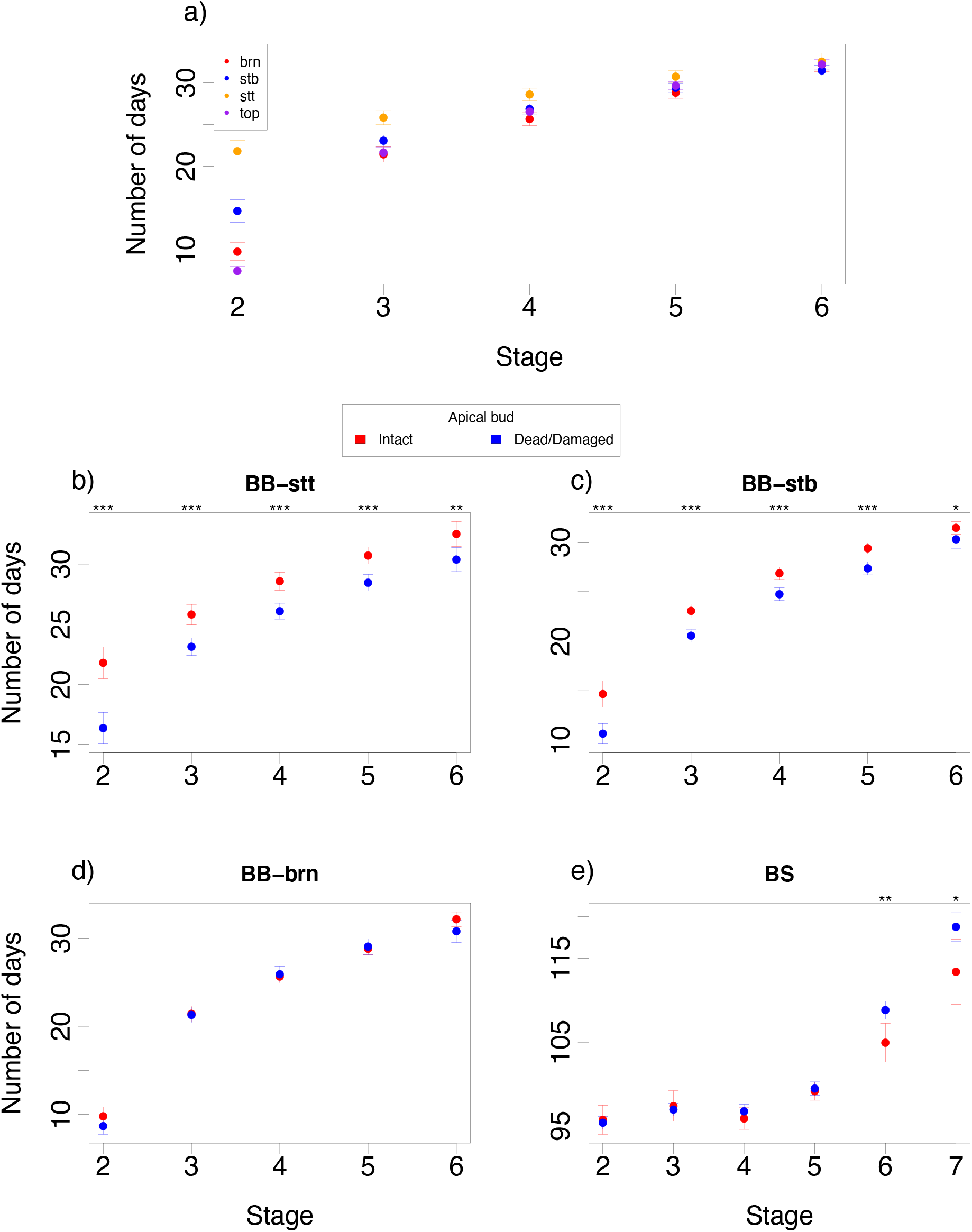
The mean of number days and 95% confidence interval to reach a stage (x-axis) for a) the four different bud types under intact apical bud, b) the stem top buds (BB-stt) under intact (red) and damaged/dead (blue) apical bud, c) the stem bottom buds (BB-stb) under intact and damaged/dead apical bud, d) the branch buds (BB-brn) under intact and damaged/dead apical bud and e) bud set (BS) under intact and damaged/dead apical bud.

### GWAS, Lindley scores, candidate genes and GO-term enrichment

The Lindley score method located between 0 and 80 significant slopes for the different traits, yielding overall 266 significant slopes (Table 1). Autumn phenology traits LS5-17 (Fig. 3a), LS2-17 and CO3-17 had the highest numbers of significant slopes with 80, 25 and 21, respectively, followed by BB4-top with 20 significant slopes. Finally, LS5-18, BB4-stb, BS2 (Fig. 3b) and BB4-brn had 14, 13, 11 and 10 significant slopes respectively. The remainder of the traits, including BB2-top (Fig. 3c), had nine or less significant slopes, with no significant slopes identified for BB2-brn (Table 1). The growth trait (DBH-17) had 4 significant slopes (Fig. 3d).

**Table 1:** Numbers of significant slopes and candidate genes (within 10 kbp of significant slope) for each of our 23 chosen autumn phenology, spring phenology and lifetime growth field and phytotron traits.

| Site | Trait | Significant slopes | Genes |
| --- | --- | --- | --- |
| Field | BB2-17 | 1 | 2 |
| Field | BB2-18 | 5 | 10 |
| Phytotron | BB2-brn | 0 | 0 |
| Phytotron | BB2-stb | 5 | 9 |
| Phytotron | BB2-stt | 5 | 22 |
| Phytotron | BB2-top | 5 | 8 |
| Field | BB4-17 | 9 | 28 |
| Field | BB4-18 | 6 | 21 |
| Phytotron | BB4-brn | 10 | 28 |
| Phytotron | BB4-stb | 13 | 30 |
| Phytotron | BB4-stt | 2 | 5 |
| Phytotron | BB4-top | 20 | 54 |
| Phytotron | BS2 | 11 | 37 |
| Phytotron | BS7 | 6 | 30 |
| Field | CO3-17 | 21 | 57 |
| Field | CO3-18 | 7 | 27 |
| Field | CO8-17 | 6 | 23 |
| Field | CO8-18 | 7 | 14 |
| Field | DBH-17 | 4 | 9 |
| Field | LS2-17 | 25 | 53 |
| Field | LS2-18 | 4 | 16 |
| Field | LS5-17 | 80 | 101 |
| Field | LS5-18 | 14 | 45 |
| Total |  | 266 | 629 |

**Figure 3:**
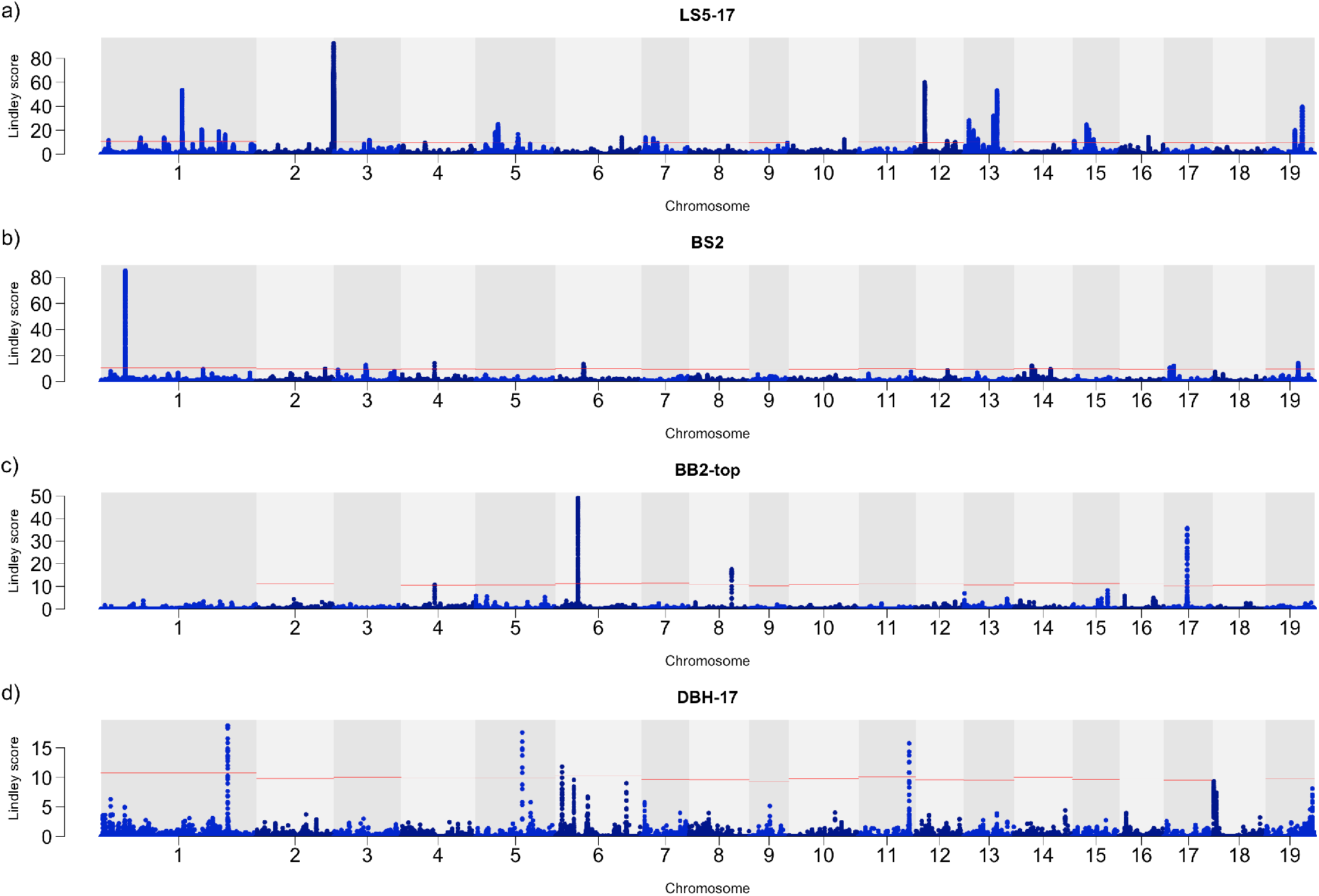
Manhattan plot of the Lindley score for a) leaf shed completion in 2017 (LS5-17), b) bud set initiation (BS2), c) bud burst initiation in the apical bud (BB2-top) and d) diameter at breast height (DBH-17). The red lines denote the chromosome-specific threshold of significance.

Between 0 and 101 genes were located within 10 kbp of the significant slopes across all traits with 629 unique genes included overall (Table 1). GO-term enrichment analysis for these genes yielded 41 enriched terms across only 8 of our traits after multiple test correction. Of these enrichments, 20 were in the category biological process, 13 in cellular component and 8 in molecular function. Overall, 5 out of the 13 significantly enriched cellular components were membranes (Table S8).

### Signatures of positive selection and GO-term enrichment

108,106 SNPs were located within 10 kbp of the significant slopes and 189 weree located in the top 0.1 percentile in the iHS selection scan (Fig. 4a). Similarly, 120,914 SNPs (higher due to the difference in handling zero recombination values) were located within 10 kbp of the significant slopes and 65 falls in the top 0.1 percentile for the H12 statistic (Fig. 4b). SNP markers surrounding candidate genes of CO3-17 and LS2-17 were significantly (p < 0.05) enriched in the top 0.1 percentile in both selection scans (Table S9) using hypergeometric distribution tests. 710 and 685 genes fell within 10 kbp of the top peaks, yielding 13 and 15 GO-term enrichments for iHS and H12 selection estimates respectively (Fig. 5, Table S8).

**Figure 4:**
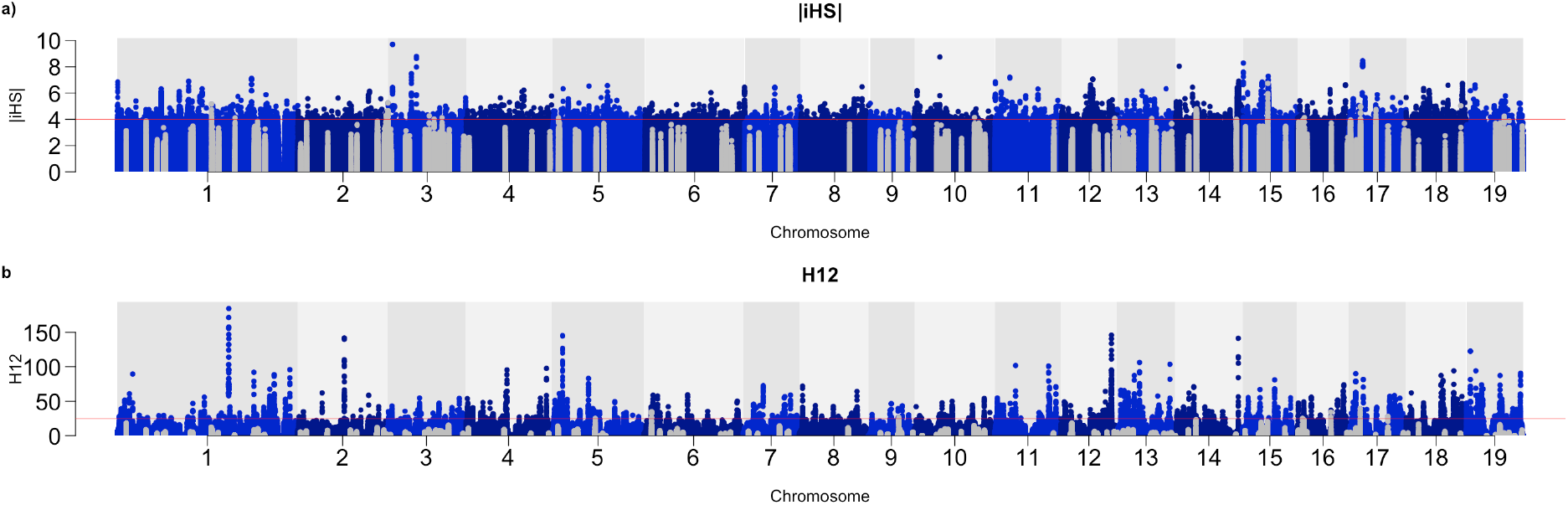
Manhattan plot of a) absolute iHS and b) H12 selection estimates. The gray dots are markers within 10 kbp of significant slopes.

**Figure 5:**
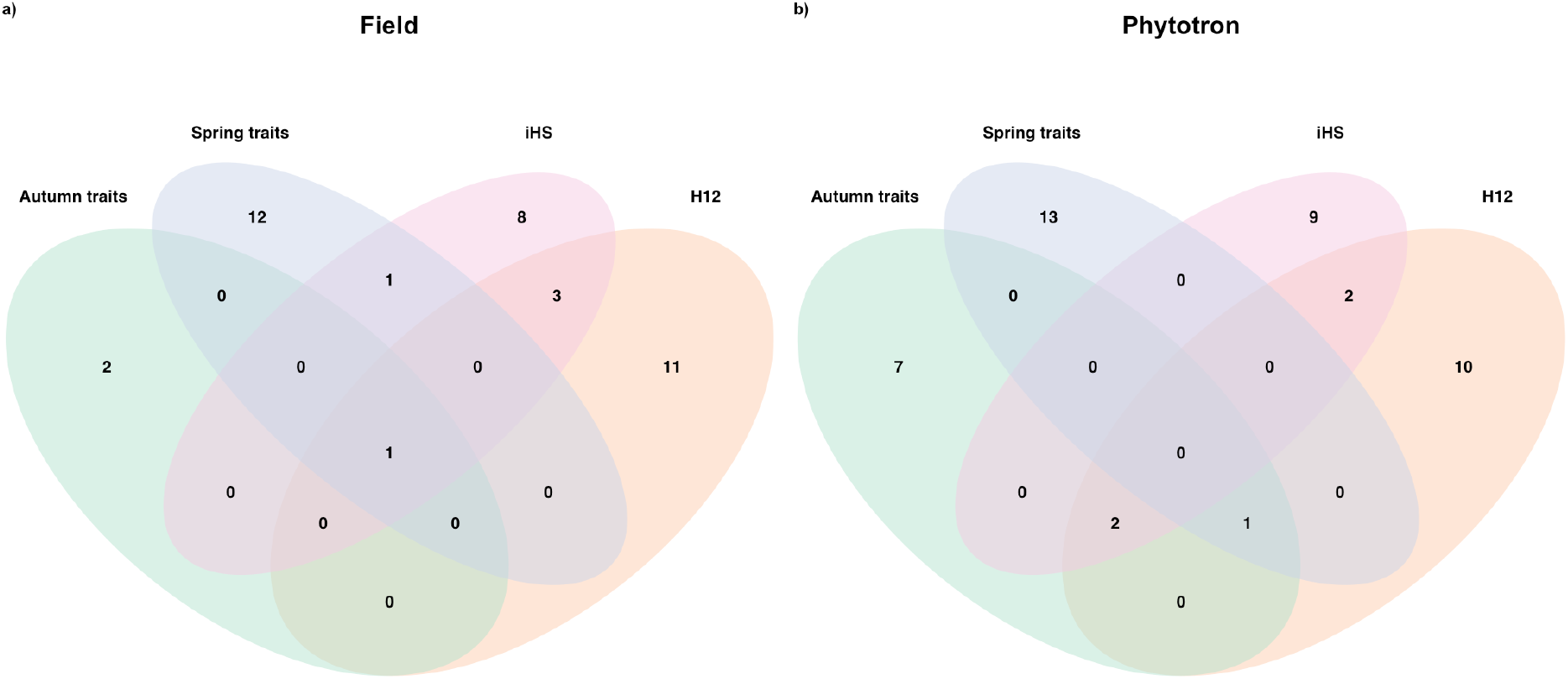
Venn diagram of enriched GO -terms for autumn and spring phenology and the two selection estimates iHS and H12 in a) field and b) phytotron.

### Autumn phenology

164 and 17 significant slopes were observed for field and phytotron autumn phenology traits and the extended slopes contained 336 and 67 candidate genes, respectively. Of the 403 candidate genes, 34 were shared between different fall phenology traits and 6 were only shared for the same trait measured in different years in the field. We observed overlaps between years for CO3 and LS2 only and no overlaps were seen for CO8 and LS5 (Fig. S4, Table S10). 23 candidate genes were shared between two different autumn phenology traits and five candidate genes were shared between four autumn phenology traits in the field. The five candidate genes were shared between CO3-17, CO3-18, LS2-17 and LS5-18 (Table S10).

Two candidate genes for autumn phenology traits were shared between our study and Evans *et al.*, (2014) and McKown *et al.*, (2014). 34 candidate genes were shared between Evans *et al.*, (2014) and our results and one candidate gene were shared between McKown *et al.*, (2014) and our results (Fig. 6a, Table S11). The AT GO-term enrichment analysis yielded 13 enrichments for autumn phenology (Fig. 5), 10 of which were found in BS2 (Fig. 5b). Four of the 13 enrichments were shared with selection estimates, 2 with H12 and 3 with iHS (Fig. 5), of which one was shared between both selection estimates and field autumn phenology traits (Fig. 5a).

**Figure 6:**
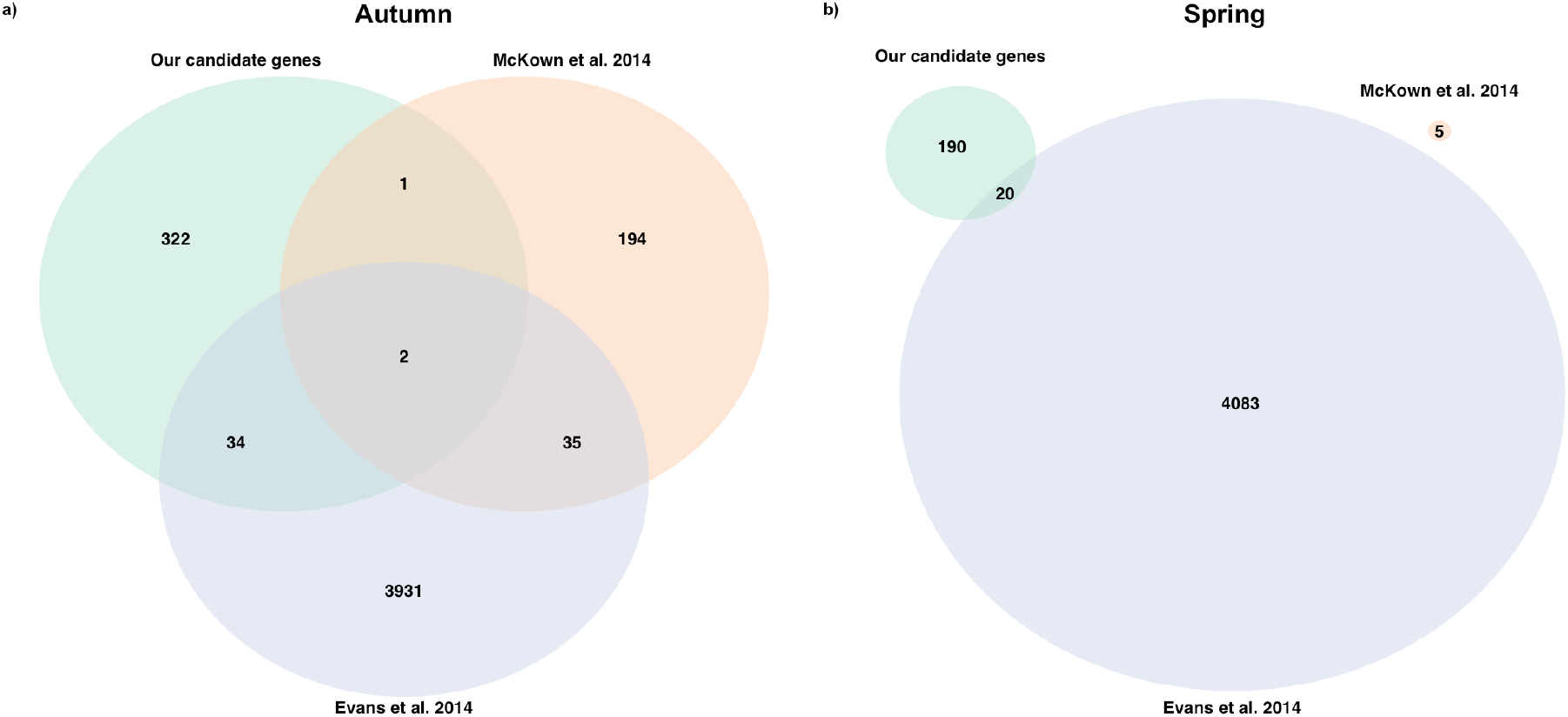
Venn diagram of overlap between our a) autumn phenology and b) spring phenology (bud burst) candidate genes and those identified in two other studies (Evans *et al.* 2014 and McKown *et al.* 2014).

### Spring phenology

In total 21 and 60 significant slopes were observed for field and phytotron spring phenology traits and the extended slopes encompass 61 and 156 candidate genes, respectively. Five genes were shared between the same trait across years in the field and 2 were shared between BB4-stb and BB4-brn in the phytotron (Fig. S4, Table S10). These 2 candidate genes constitute the only overlap for the phytotron bud burst as the four phytotron bud burst traits showed no within stage overlap in candidate genes for either stage, BB2 or BB4 (§). No overlap was observed between the phytotron and field estimates for these stages either (Table S10). Additionally, a total of 7 candidate genes were shared between at least one autumn and one spring phenology trait.

For spring phenology traits, we observed no candidate genes that were shared between our results and McKown *et al.*, (2014), while a total of 20 genes were shared between our results and Evans *et al.*, (2014) (bud burst) (Fig. 6b, Table S11). There were 28 enrichments found for spring phenology traits in the AT GO-term analysis (Fig. 5). Finally, 2 GO -term enrichments were shared between spring phenology and H12 and iHS selection estimates respectively (Fig. 5).

### Phenology candidate genes for adaptation to northern Europe

Acyl transferase/acyl hydrolase/lysophospholipase superfamily protein (Potri.001G209500) with function in lipid metabolism and auxin response factor 1 (Potri.004G228800) were shared between years in BB2 and CO3 respectively. Both BB4 (Potri.016G030500, Potri.016G030600) and LS2 (Potri.016G033500, Potri.016G033600, Potri.016G033700) had DNA/RNA helicase protein shared between the years. Two Prolyl oligopeptidase family protein coding genes were shared between the four traits CO3-17, CO3-18, LS2-17 and LS5-18. The genes shared between CO and LS in year 2018 included genes such as xyloglucan endotransglucosylase/hydrolase 5 (Potri.003G159700) with function in primary cell wall metabolism (Hyodo *et al.*, 2003) and ENTH/VHS/GAT protein (Potri.002G038000) with roles in membrane trafficking and cytokinesis (De Craene *et al.*, 2012).

## Discussion

### Phenotypic variation and heritability

All of the traits had appreciable levels of phenotypic variation, with BB2-top and BS2 being the least variable traits (Fig. 1e & f). Similarly, the vast majority of traits had appreciable levels of heritability (Table S7). The clear exceptions being BB2-top and BS2 which showed a general lack of variability and also low heritabilites (Fig. 1e & f, Table S7). Heritabilities are known to vary between different environments, though for most species and traits, measurements taken in a controlled environment usually produce higher heritability estimates (Ali & Johnson, 1999).

### Genetic architecture of phenology and growth

We identified a total 266 significant slopes across the 23 study traits. Among the traits, LS5-17 has more than three times as many significant slopes than the trait with next highest number of slopes (Fig. 3a, Table 1). This supports earlier evidence for the complex genetic basis of autumn phenology traits in *P. trichocarpa* (Evans *et al.*, 2014; McKown *et al.*, 2014). For growth (DBH-17) we only identified 4 significant slopes (Fig. 3d, Table 1), likely due to the highly polygenic nature of growth traits and the low power to detect loci with only small effects even with modern GWAS methods (Du *et al.*, 2016; Müller *et al.*, 2019).

We observed an earlier initiation and completion of both autumn and spring phenology transitions in 2018 compared to 2017 (Fig. 1). We also identified noticeably fewer significant slopes for autumn phenology traits in 2018 compared to 2017 (Table 1). Bud burst (Nicotra *et al.*, 2010; Tansey *et al.*, 2017), leaf shed and leaf coloring (Ghelardini *et al.*, 2014) have all been found to be affected by weather conditions, which likely explains many of the observed differences between the years. In 2017 the average temperature was above 5 °C from May to October with an average temperature of 12.8 °C whereas the average temperature in 2018 was above 5 °C already in April and lasted until October with an average temperature of 13.9 °C. The difference in mean temperature is largely caused by an early and very warm spring and an exceptional heatwave during July in 2018. Precipitation was also substantially lower 2018 compared to 2017 (Table S2).

Changes in lipid and protein metabolism have been previously observed during phenology transitions in *Populus* and other tree species (Hoffman *et al.*, 2010; Lee *et al.*, 2014). Lipid contents of various membranes are well-established indicators of cell status, such as cold hardiness (Kedrowski, 1980; Lee *et al.*, 2014). Our GO-term analyses support this as we uncovered 5 enrichments across our traits that are directly linked to membrane structures three of which were in GO-term plasma membrane (GO:0005886) found in BB-18, BS and CO-17 (Table S8).

### Signatures of positive selection and GO-term enrichment

Positive selection often drives local adaptation (Fournier-Level *et al.*, 2011), leaving detectable signatures in genetic variation across the genome. Using two test statistics, iHS (Voight *et al.*, 2006) and H12 (Garud *et al.*, 2015), we identified clear signatures of positive selection at multiple locations across the genomes of our *P. trichocarpa* study population. We observed hundreds of markers within 10 kbp of significant slopes in the top 0.1 percentile of estimated selection values for both of the selection scans (Fig. 4). Thus, many of the significant slopes we identify likely also correspond to genome regions that have been under positive selection in the native environments of the parents of our study population.

The GO-term analysis of putative candidate genes under selection also yielded enrichments in plasma membrane (GO:0005886) for both statistics, chloroplast envelope (GO:0009941) for iHS and membrane (GO:0016020) for H12, lending further support to the importance of membrane structures for adaptation to northern climates (Kedrowski, 1980; Lee *et al.*, 2014). Among other noteworthy enrichments was response to cold (GO:0009409) enriched for iHS (Table S8) intuitively linking to both autumn and spring phenology as some aspects of both have been found to be temperature dependent in *Populus* (Fracheboud *et al.*, 2009).

### Autumn phenology

Previous work has shown that bud set initiation is genetically more well-defined than bud set completion in *Populus* (Fracheboud *et al.*, 2009; Rohde *et al.*, 2011). Our findings support this, as we observe almost twice as many significant slopes for BS2 than BS7 in the phytotron. Initiation of both bud set and leaf senescence has been previously observed to be consistent between years and conditions for the same trees, suggesting a more stringent genetic control of the initiation of autumn phenology traits (Fracheboud *et al.*, 2009; Rohde *et al.*, 2011; Richards *et al.*, 2020). Our results are in line with this, as the 11 candidate genes we found shared between years were exclusively found in the early stages (CO3 and LS2) (Fig. S4, Table S10). Similarly, autumn phenology traits have been previously shown to have a degree of shared genetic architecture (McKown *et al.*, 2014) and be genetically correlated in this population (Richards *et al.*, 2020). Our results support this as we observe a notable overlap in candidate genes between different autumn phenology traits in the field including shared associations of five candidate genes across four field traits. The largest of these overlaps occurring between CO3-18 and LS5-18 and encompassing 15 candidate genes (Table S10), could potentially be taken as evidence for a systematic stress response to heat and/or drought (Table S10, Table S2). The senescence hastening effects of these stresses have been observed in model species (Munné-Bosch & Alegre, 2004; Woo *et al.*, 2019) fitting well with our observations in 2018.

We identified two candidate genes for autumn phenology traits that were also identified in the earlier studies of Evans *et al.*, (2014) and McKown *et al.*, (2014) (Fig. 6a). These genes were glucan synthase-like 12 (Potri.003G214200) shared between LS2-18 and bud set in both Evans *et al.*, (2014) and McKown *et al.*, (2014), and additional traits such as yellowing and leaf drop in the latter, and glucuronidase 2 (Potri.015G049100) shared between LS5-17 and leaf drop in McKown *et al.*, (2014) and bud set in Evans *et al.*, (2014). As both of these have functions in metabolism of complex carbohydrates (https://uniprot.org), they may have roles in cell wall degradation or production of storage carbohydrates. An additional 34 candidate genes that we identify in this study are also shared with Evans *et al.*, (2014) but only a single gene is shared with McKown *et al.*, (2014) (Fig. 6a). These overlaps are similar in magnitude to previous comparisons (Zhang *et al.*, 2019) and suggest that the genetic architecture of autumn phenology traits is shared across both native and novel environments. However, the relatively large numbers of study specific candidate genes also hint at the complexity of these traits under the variable natural conditions.

Both the selection statistics showed significant enrichment with GWAS hits in the top 0.1 percentile for CO3-17 and LS2-17 (Table S9). This suggests that the start of the autumn phenology transition is a more important adaptation than the completion, a view that has been supported in earlier studies in *Populus* (Fracheboud *et al.*, 2009). Furthermore, the fact that we observe significant enrichments in these two traits but fail to detect any enrichments in the corresponding traits in 2018 lends support to our view that different environmental factors were driving autumn senescence across the two years. These observations suggest that results from the more usual year of 2017 and in particular the candidate genes we identify, are more relevant for driving local adaptation.

### Spring phenology

The apical meristem has more functions than axillary meristems in species with apical dominance. One such extra function is the inhibition of axillary buds through apically produced auxins (Chatfield *et al.*, 2000; Qiu *et al.*, 2019) that are basipetally transported to the buds along the stem (Ljung *et al.*, 2001). We studied the timing of bud burst of different buds in our phytotron experiment and the results reflect the apical dominance effect, as the apical bud (top) initiates bud burst earlier than other types of buds (Fig. 2a). Furthermore, the stem top 50% (stt) and stem bottom 50% (stb) buds display considerably different timing of bud burst conditional on the presence of a functional apical bud (Fig. 2b & c).

The higher numbers of significant genes observed in phytotron in combination with a general lack of overlap in candidate genes between bud burst traits supports not only the well-established view that bud burst is a highly plastic trait but also that is has a complex genetic basis (Evans *et al.*, 2014; McKown *et al.*, 2014). The lack of overlap in candidate genes identified between the bud burst traits in the phytotron experiment suggests that the genes controlling bud burst in different parts of the plant are unique (Fig. 7), which may have an effect on the comparability of results between studies.

**Figure 7:**
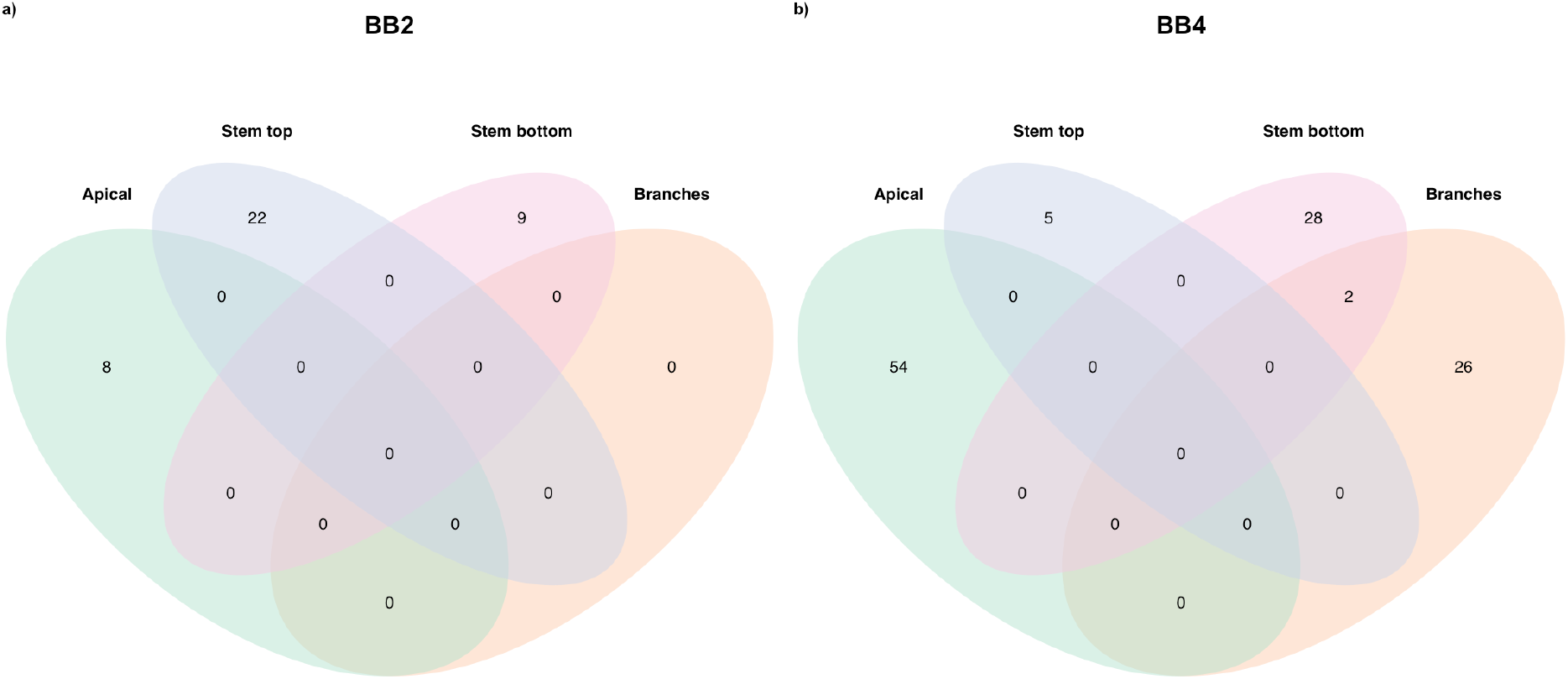
Venn diagram of the four different bud types in phytotron for a) the initiation stage (BB2) and b) for the completion stage (BB4).

A total of 5 candidate genes were identified for bud burst, with 2 and 3 shared between the initiation stage BB2 and the completion stage BB4 respectively. This aligns nicely with the genetic correlations found previously (Richards *et al.*, 2020), but somewhat surprisingly suggests a more consistent genetic control in bud burst between years than in autumn phenology (Fig. S4). Comparing our results to Evans *et al.*, (2014), we observe an overlap of 20 genes candidate genes, suggesting that there is a degree of similarity in the genetic control of bud burst between the native and the novel environments. However, this overlap is substantially lower that what we observed for the autumn phenology traits (Fig. 6a & b).

### Phenology candidate genes for adaptation to northern Europe

Candidate genes identified for phenology traits across the two years offer potential targets for adaptive improvement of *P. trichocarpa* to northern European conditions. We observed candidate genes with functions in auxin metabolism, lipid metabolism and as helicases (Table S10). Auxins have a well-established role in delaying of senescence in plants (Mueller-Roeber & Balazadeh, 2014; Woo *et al.*, 2019). Helicases have function in separating DNA strands likely contributing to the large changes in transcriptome phenology transitions constitute in *Populus trichocarpa* (Howe *et al.*, 2015). Shared between the four autumn phenology traits of CO3-17, CO3-18, LS2-17 and LS5-18, oligopeptidases function in protein catabolism which is in line with the observations that important proteins such as chlorophyll and carotenoids are broken down during leaf senescence (Kmiec *et al.*, 2013; Poret *et al.*, 2016). As the summer of 2018 was extremely warm, the high number of genes shared between CO and LS could offer insight into stress induced senescence. The genes shared between the traits would seem to be in consensus with previously established roles of the cell wall and cytokines in phenology transitions (Howe *et al.*, 2015; Woo *et al.*, 2019).

## Conclusions

The study presented here is the first to study the genetic basis of phenology traits in a population of *P. trichocarpa* introduced to northern Europe. We find complex genetic architectures underlying all phenology and growth traits, and identify multiple putative candidate genes. Together with results from Richards *et al.*, (2020) this means there is potential for adaptive improvement in our *P. trichocarpa* population. Many of the candidate genes we identify function in cell membranes or cell wall, which both have significant biological functions during phenology transitions in the novel environment of northern Europe. Comparison of candidate genes with studies performed in the native range show some overlap for both autumn and spring phenology transitions although the latter show far less overlap. Coupling these observations with the evidence for significant enrichment of SNPs under selection with regions harboring loci controlling autumn phenology transitions, this agrees with earlier observations of a more genetically well-defined control of the initiation of autumn phenology compared to both the completion of autumn phenology transitions and spring phenology in general.

## Acknowledgements

This project was funded by the Swedish Research Council Formas as part of the Climate-Adapted Poplar (CLAP) project (project number 942-2016-1). The authors acknowledge support from Science for Life Laboratory and the National Genomics Infrastructure (NGI for providing assistance with massively parallel sequencing (projects P7512 and P11086). Access to computational infrastructure was provided by the Swedish National Infrastructure for Computing (SNIC) through Uppsala Multidisciplinary Center for Advanced Computational Science (UPPMAX) under projects SNIC 2017/7-291, SNIC 2018/3-552, SNIC 2019/3-597 and sllstore2017050.

## Author contributions

ACW and PKI designed the research, RAP, TR, AK and ACW collected all field and phytotron data, RAP, TR, MR and PKI performed all data analyses, RAP and PKI wrote the manuscript, RAP, TR, MRT, AK, ACW and PKI edited the manuscript.

## Data availability

The raw sequencing reads for all individuals included in the study are available from ENA under study number PRJEB38910 (https://www.ebi.ac.uk/ena/data/view/PRJEB38910). All phenotype data are available from the authors upon request. Additional scripts and files used for all analyses are available at https://github.com/parkingvarsson/CLAP.

