## Supplementary Figures and Tables for "The genetic basis of adaptation in phenology in an introduced population of Black Cottonwood (*Populus trichocarpa*, Torr. & Gray)"

Table S1: Schematic of crosses and numbers of individuals produced by a cross between the parent individuals. Mothers are on the left (yellow) and fathers on top (blue). The parents in bold and all offspring are present in the Krusenberg field trial.

| | | Region | Alaska | | British Columbia | Idaho | Oregon | Washington x Oregon | Washington | Open Pollination, Koster, Holland | | |
|---|---|---|---|---|---|---|---|---|---|---|---|---|
| | | Latitude | 58,30 (15) | 60,00 | 49,15 | 48,25 | 45,35 | | 45,50 (75) | | | |
| Region | Latitude | Clone | **293** | **278** | **193** | 214 | 159 | 236 | **299** | 187 | 209 | 290 |
| Alaska | 58,30 | **216** | | | 5 | | 1 | | 5 | | | 5 |
| | 59,20 (500) | **291** | | | | | | 2 | | 1 | | |
| | 60,37 (400) | **265** | 3 | 3 | | 1 | | | 2 | 2 | | |
| British Columbia | 49,00 | **230** | 7 | 4 | | 3 | | 2 | 6 | | | |
| | | 96 | | | | | | | 1 | | | |
| Idaho x Oregon | | 237 | 2 | 1 | | 3 | | | 3 | 1 | | 3 |
| | | **240** | | 4 | | | | 1 | 21 | | 2 | |
| Oregon | 44,30 | 271 | 1 | 2 | | | | 1 | 4 | | | 1 |
| | | 301 | | | | | | | | 2 | | 1 |

Table S2: Monthly mean temperature and rainfall information for years 2017 and 2018.

| | 2017 | | 2018 | |
|---|---|---|---|---|
| | Temp. (°C) | Rainfall (mm) | Temp. (°C) | Rainfall (mm) |
| Jan | -1.6 | 15.8 | -1.4 | 53.6 |
| Feb | -1.14 | 19.2 | -4.56 | 19.1 |
| Mar | 2.25 | 28.5 | -3.11 | 23.1 |
| Apr | 3.76 | 26.2 | 6.05 | 36.3 |
| May | 10.45 | 10.4 | 15.33 | 6.7 |
| Jun | 14.7 | 50.6 | 16.3 | 20.7 |
| Jul | 16.62 | 14.6 | 21.6 | 81.7 |
| Aug | 15.78 | 61.1 | 17.77 | 68.7 |
| Sep | 12.16 | 68.2 | 12.86 | 42.2 |
| Oct | 6.79 | 89.9 | 7.48 | 21.1 |
| Nov | 2.61 | 70.5 | 3.41 | 24.3 |
| Dec | 0.18 | 49.9 | -0.28 | 31.1 |

Table S3: Drawing, picture and description of all 6 stages of bud burst. Drawings reproduced on permission of the owner Alfas Pliura of Lithuanian Research Centre for Agriculture and Forestry.

| Drawing of stage | **Picture of stage** | **Description of stage** |
|---|---|---|
| 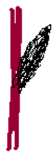 | 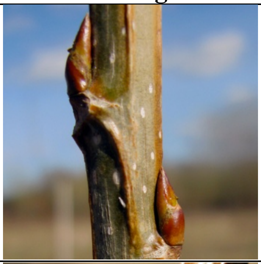 | **Stage 1**. Bud swollen, no initials visible. Stage 1,5 if ca 1 mm initial tip visible while bud scales are completely closed |
| 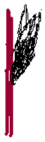 | 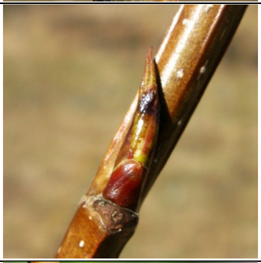 | **Stage 2**. Buds are opening, initials visible protruding up to one length of bad scales. |
| NA | 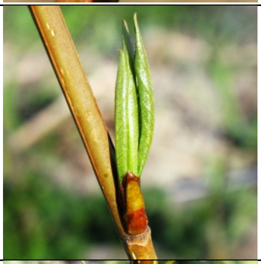 | **Stage 3**. Buds are still opening, the leaf primordia elongated, rolled up and longer than bud scales |
| 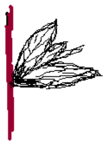 | 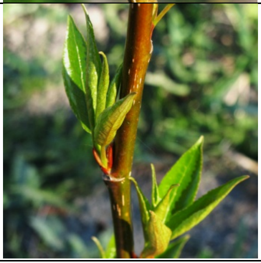 | **Stage 4.** Leaves half-shed, bad scales falling off. Stage 4,5 if most of the leaves but not all are shed. |

| | | |
|---|---|---|
| 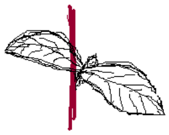 | NA | **Stage 5.** Leaves completely shed |
| 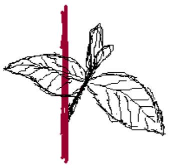 | NA | **Stage 6.** >1 cm shoot increment, leaves completely shed |

Table S4: Autumn coloration scoring scheme for 2017-2018. Original 2017 and 2018 describe the original scoring systems used in the corresponding years. converted 2017 shows, which stages the original 2017 scores correspond in the 2018 scoring system.

| Percentage of yellowing | Original 2017 | Original 2018 | Converted 2017 |
|---|---|---|---|
| 0% | 1 | 1 | 1 |
| | | 2 | |
| 25% | 2 | 3 | 3 |
| | | 4 | |
| 50% | 3 | 5 | 5 |
| 75% | 4 | 6 | 6.5 |
| | | 7 | |
| 100% | 5 | 8 | 8 |

Table S5: Phytotron simulated seasons temperature, light conditions and humidity information.

| Date | Temperature day (ºC) | Temperature night (ºC) | Day length (h) | Light intensity (umol) | Humidity (%) |
|---|---|---|---|---|---|
| 20.03.2018 | 22 | 18 | 18 | 350 | 80 |
| 24.04.2018 | 22 | 18 | 8 | 100 | 80 |
| 01.05.2018 | 4 | 4 | 8 | 100 | 90 |
| 10.09.2018 | 10 | 10 | 12 | | |
| 20.09.2018 | 15 | 15 | 15 | | |
| 30.09.2018 | 17 | 17 | 18 | | |
| 10.10.2018 | 16 | 16 | 17 | | |
| 20.10.2018 | 11 | 11 | 15 | | |
| 30.10.2018 | 7.5 | 7.5 | 12 | | |
| 09.11.2018 | 4 | 4 | 10 | | |
| 19.11.2018 | 4 | 4 | 7 | | |
| 29.11.2018 | 4 | 4 | 6 | | |
| 09.12.2018 | 4 | 4 | 7 | | |
| 19.12.2018 | 4 | 4 | 10 | | |

Table S6: Sequencing breadth and depth coverage of the 121 sequenced individuals. Values have been rounded to three decimals.

| SAMPLE | DEPTH | BREADTH |
|---|---|---|
| P11086_101 | 19.281 | 0.897 |
| P11086_102 | 19.59 | 0.902 |
| P11086_103 | 19.167 | 0.899 |
| P11086_104 | 15.197 | 0.894 |
| P11086_105 | 16.726 | 0.895 |
| P7512_101 | 30.848 | 0.911 |
| P7512_102 | 22.041 | 0.907 |
| P7512_103 | 25.771 | 0.907 |
| P7512_104 | 19.793 | 0.906 |
| P7512_105 | 20.279 | 0.903 |
| P7512_106 | 24.214 | 0.896 |
| P7512_107 | 27.784 | 0.908 |
| P7512_109 | 21.203 | 0.902 |
| P7512_110 | 12.205 | 0.884 |
| P7512_111 | 20.394 | 0.906 |
| P7512_112 | 22.627 | 0.9 |
| P7512_113 | 29.538 | 0.907 |
| P7512_114 | 24.842 | 0.911 |
| P7512_115 | 25.263 | 0.907 |
| P7512_116 | 18.111 | 0.906 |
| P7512_117 | 20.471 | 0.9 |
| P7512_118 | 24.39 | 0.904 |
| P7512_119 | 12.887 | 0.894 |
| P7512_120 | 25.466 | 0.906 |
| P7512_121 | 30.1 | 0.906 |
| P7512_122 | 18.896 | 0.903 |
| P7512_123 | 22.537 | 0.906 |
| P7512_124 | 12.746 | 0.895 |
| P7512_125 | 19.917 | 0.901 |
| P7512_126 | 19.027 | 0.902 |
| P7512_127 | 23.981 | 0.908 |
| P7512_128 | 17.109 | 0.899 |
| P7512_129 | 17.024 | 0.902 |
| P7512_131 | 18.485 | 0.901 |
| P7512_132 | 21.5 | 0.904 |
| P7512_133 | 13.617 | 0.898 |
| P7512_134 | 12.272 | 0.895 |
| P7512_135 | 21.195 | 0.903 |
| P7512_136 | 19.323 | 0.902 |
| P7512_138 | 23.152 | 0.907 |
| P7512_139 | 17.866 | 0.902 |

| | | |
|---|---|---|
| P7512_140 | 15.47 | 0.898 |
| P7512_141 | 21.393 | 0.906 |
| P7512_142 | 24.329 | 0.904 |
| P7512_143 | 18.81 | 0.903 |
| P7512_144 | 19.342 | 0.9 |
| P7512_145 | 25.089 | 0.909 |
| P7512_146 | 44.43 | 0.918 |
| P7512_147 | 23.154 | 0.903 |
| P7512_148 | 21.279 | 0.909 |
| P7512_149 | 20.692 | 0.908 |
| P7512_150 | 19.929 | 0.905 |
| P7512_151 | 21.089 | 0.904 |
| P7512_152 | 20.233 | 0.903 |
| P7512_153 | 21.629 | 0.906 |
| P7512_154 | 26.59 | 0.908 |
| P7512_155 | 25.857 | 0.909 |
| P7512_156 | 23.673 | 0.908 |
| P7512_157 | 28.413 | 0.91 |
| P7512_158 | 24.345 | 0.912 |
| P7512_159 | 17.221 | 0.902 |
| P7512_160 | 18.091 | 0.902 |
| P7512_161 | 17.535 | 0.903 |
| P7512_162 | 29.995 | 0.917 |
| P7512_163 | 36.829 | 0.917 |
| P7512_164 | 26.973 | 0.912 |
| P7512_165 | 31.253 | 0.912 |
| P7512_166 | 18.89 | 0.905 |
| P7512_168 | 14.342 | 0.899 |
| P7512_169 | 21.486 | 0.904 |
| P7512_170 | 47.833 | 0.917 |
| P7512_171 | 18.886 | 0.903 |
| P7512_172 | 21.603 | 0.907 |
| P7512_173 | 21.431 | 0.908 |
| P7512_174 | 38.077 | 0.917 |
| P7512_175 | 23.981 | 0.909 |
| P7512_176 | 21.224 | 0.907 |
| P7512_177 | 30.228 | 0.919 |
| P7512_178 | 25.63 | 0.919 |
| P7512_179 | 30.368 | 0.915 |
| P7512_180 | 22.934 | 0.905 |
| P7512_181 | 15.927 | 0.897 |
| P7512_182 | 23.244 | 0.906 |

| | | |
|---|---|---|
| P7512_184 | 31.17 | 0.91 |
| P7512_185 | 32.415 | 0.911 |
| P7512_186 | 30.165 | 0.906 |
| P7512_187 | 45.819 | 0.916 |
| P7512_188 | 37.223 | 0.912 |
| P7512_189 | 17.618 | 0.901 |
| P7512_190 | 19.131 | 0.9 |
| P7512_191 | 28.347 | 0.906 |
| P7512_192 | 28.183 | 0.905 |
| P7512_193 | 26.48 | 0.908 |
| P7512_194 | 46.832 | 0.915 |
| P7512_195 | 32.954 | 0.911 |
| P7512_196 | 15.131 | 0.898 |
| P7512_201 | 30.811 | 0.911 |
| P7512_202 | 14.874 | 0.899 |
| P7512_204 | 33.022 | 0.912 |
| P7512_205 | 13.249 | 0.898 |
| P7512_206 | 41.296 | 0.917 |
| P7512_208 | 31.745 | 0.912 |
| P7512_209 | 15.72 | 0.898 |
| P7512_210 | 49.304 | 0.917 |
| P7512_211 | 46.784 | 0.913 |
| P7512_212 | 48.918 | 0.922 |
| P7512_213 | 21.381 | 0.906 |
| P7512_214 | 32.925 | 0.916 |
| P7512_215 | 15.937 | 0.894 |
| P7512_216 | 31.262 | 0.901 |
| P7512_217 | 24.773 | 0.872 |
| P7512_219 | 17.099 | 0.85 |
| P7512_220 | 17.526 | 0.902 |
| P7512_221 | 43.977 | 0.914 |
| P7512_222 | 21.007 | 0.907 |
| P7512_223 | 16.213 | 0.899 |
| P7512_225 | 37.838 | 0.922 |
| P7512_226 | 17.068 | 0.897 |
| P7512_227 | 15.671 | 0.897 |
| P7512_228 | 13.816 | 0.892 |
| P7512_229 | 23.007 | 0.909 |

Table S7: Heritabilites of chosen traits.

| Trait | Narrow sense heritability | Number of genotypes |
|---|---|---|
| DBH-17 | 0.745 | 109 |
| BB2-17 | 0.698 | 109 |

| | | |
|---|---|---|
| BB4-17 | 0.777 | 108 |
| CO3-17 | 0.593 | 104 |
| CO8-17 | 0.839 | 107 |
| LS2-17 | 0.464 | 108 |
| LS5-17 | 0.609 | 86 |
| BB2-18 | 0.567 | 109 |
| BB4-18 | 0.529 | 108 |
| CO3-18 | 0.474 | 108 |
| CO8-18 | 0.665 | 102 |
| LS2-18 | 0.647 | 109 |
| LS5-18 | 0.880 | 105 |
| BB2-brn | 0.160 | 67 |
| BB4-brn | 0.529 | 69 |
| BB2-stb | 0.171 | 93 |
| BB4-stb | 0.755 | 92 |
| BB2-stt | 0.581 | 93 |
| BB4-stt | 0.824 | 89 |
| BB2-top | 0.028 | 83 |
| BB4-top | 0.391 | 79 |
| BS2 | 0.107 | 51 |
| BS7 | 0.555 | 82 |

Table S8: GO -term enrichment analysis results.

| Trait | GO type | **GO id** | **Pval (COrrected)** | **Statistics** | **Description** |
|---|---|---|---|---|---|
| BB-17 | Molecular function | GO:0009055 | 1.747e-03 | 3/7 \| 502/24493 | electron carrier activity |
| BB-17 | Cellular Component | GO:0005783 | 1.768e-02 | 2/5 \| 354/23421 | endoplasmic reticulum |
| BB-17 | Molecular function | GO:0020037 | 9.081e-04 | 3/7 \| 373/24493 | heme binding |
| BB-17 | Molecular function | GO:0005506 | 1.136e-03 | 3/7 \| 365/24493 | iron ion binding |
| BB-17 | Molecular function | GO:0004497 | 1.133e-03 | 3/7 \| 252/24493 | monooxygenase activity |
| BB-17 | Biological process | GO:0055114 | 3.888e-02 | 3/7 \| 1403/24222 | oxidation-reduction process |
| BB-17 | Molecular function | GO:0016705 | 7.032e-04 | 3/7 \| 271/24493 | oxidoreductase activity, acting on paired donors, with inCOrporation or reduction of molecular oxygen |
| BB-17 | Molecular function | GO:0019825 | 7.045e-03 | 2/7 \| 232/24493 | oxygen binding |
| BB-18 | Biological process | GO:0009926 | 3.199e-02 | 3/85 \| 42/24222 | auxin polar transport |
| BB-18 | Biological process | GO:0042742 | 1.084e-02 | 6/85 \| 176/24222 | defense response to bacterium |
| BB-18 | Cellular Component | GO:0010008 | 2.553e-03 | 2/86 \| 3/23421 | endosome membrane |
| BB-18 | Biological process | GO:0048527 | 1.261e-02 | 3/85 \| 28/24222 | lateral root development |
| BB-18 | Biological process | GO:0009808 | 1.072e-02 | 2/85 \| 4/24222 | lignin metabolic process |

| | | | | | |
|---|---|---|---|---|---|
| BB-18 | Cellular Component | GO:0005886 | 1.881e-03 | 19/86 \| 1929/23421 | plasma membrane |
| BB-brn | Biological process | GO:0048364 | 2.290e-02 | 2/17 \| 99/24222 | root development |
| BB-brn | Biological process | GO:0006812 | 2.471e-02 | 2/17 \| 108/24222 | cation transport |
| BB-brn | Biological process | GO:0009611 | 3.139e-02 | 2/17 \| 143/24222 | response to wounding |
| BB-brn | Biological process | GO:0008152 | 3.322e-02 | 4/17 \| 1033/24222 | metabolic process |
| BB-brn | Biological process | GO:0006351 | 3.796e-02 | 2/17 \| 90/24222 | transcription, DNA-dependent |
| BB-brn | Biological process | GO:0009058 | 4.281e-02 | 2/17 \| 179/24222 | biosynthetic process |
| BB-brn | Cellular Component | GO:0008194 | 1.637e-03 | 3/16 \| 100/24493 | UDP-glyCOsyltransferase activity |
| BB-brn | Cellular Component | GO:0016758 | 5.753e-03 | 3/16 \| 193/24493 | transferase activity, transferring hexosyl groups |
| BB-brn | Cellular Component | GO:0016779 | 6.519e-03 | 2/16 \| 47/24493 | nucleotidyltransferase activity |
| BB-brn | Cellular Component | GO:0016757 | 9.046e-03 | 3/16 \| 347/24493 | transferase activity, transferring glyCOsyl groups |
| BB-stb | Biological process | GO:0040007 | 1.314e-02 | 2/17 \| 21/24222 | growth |
| BB-stb | Biological process | GO:0009414 | 1.995e-02 | 3/17 \| 189/24222 | response to water deprivation |
| BB-stb | Biological process | GO:0042538 | 2.424e-02 | 2/17 \| 49/24222 | hyperosmotic salinity response |
| BB-stt | Molecular function | GO:0035251 | 3.488e-02 | 2/39 \| 23/24493 | UDP-gluCOsyltransferase activity |
| BS | Biological process | GO:0010119 | 2.671e-03 | 4/108 \| 28/24222 | regulation of stomatal movement |
| BS | Biological process | GO:0010227 | 1.061e-02 | 3/108 \| 17/24222 | floral organ aBScission |
| BS | Biological process | GO:0009414 | 1.955e-02 | 6/108 \| 189/24222 | response to water deprivation |
| BS | Biological process | GO:0030007 | 2.467e-02 | 2/108 \| 5/24222 | cellular potassium ion homeostasis |
| BS | Biological process | GO:0030833 | 4.108e-02 | 2/108 \| 8/24222 | regulation of actin filament polymerization |
| BS | Biological process | GO:0009416 | 4.418e-02 | 5/108 \| 157/24222 | response to light stimulus |
| BS | Cellular Component | GO:0005634 | 5.239e-03 | 22/98 \| 2161/23421 | nucleus |
| BS | Cellular Component | GO:0005885 | 1.163e-02 | 2/98 \| 9/23421 | Arp2/3 protein COmplex |
| BS | Cellular Component | GO:0000326 | 1.364e-02 | 2/98 \| 7/23421 | protein storage vacuole |
| BS | Cellular Component | GO:0005886 | 1.459e-02 | 19/98 \| 1929/23421 | plasma membrane |
| CO-17 | Cellular Component | GO:0005886 | 2.346e-02 | 10/37 \| 1929/23421 | plasma membrane |
| H12 | Cellular Component | GO:0009570 | 1.325e-02 | 11/182 \| 396/23421 | chloroplast stroma |
| H12 | Cellular Component | GO:0005737 | 2.879e-02 | 14/182 \| 703/23421 | cytoplasm |
| H12 | Cellular Component | GO:0005829 | 4.151e-02 | 10/182 \| 465/23421 | cytosol |

| | | | | | |
|---|---|---|---|---|---|
| H12 | Biological process | GO:0009793 | 2.472e-02 | 11/196 \| 335/24222 | embryo development ending in seed dormancy |
| H12 | Biological process | GO:0009558 | 4.726e-02 | 2/196 \| 5/24222 | embryo sac cellularization |
| H12 | Cellular Component | GO:0005615 | 3.880e-02 | 2/182 \| 9/23421 | extracellular space |
| H12 | Cellular Component | GO:0016020 | 4.860e-02 | 27/182 \| 2001/23421 | membrane |
| H12 | Biological process | GO:0032876 | 4.726e-02 | 2/196 \| 5/24222 | negative regulation of DNA endoreduplication |
| H12 | Cellular Component | GO:0005634 | 3.680e-04 | 37/182 \| 2161/23421 | nucleus |
| H12 | Biological process | GO:0009626 | 2.435e-02 | 4/196 \| 33/24222 | plant-type hypersensitive response |
| H12 | Cellular Component | GO:0005886 | 1.022e-02 | 30/182 \| 1929/23421 | plasma membrane |
| H12 | Biological process | GO:0006355 | 4.714e-02 | 23/196 \| 1388/24222 | regulation of transcription, DNA-dependent |
| H12 | Biological process | GO:0010583 | 2.283e-02 | 6/196 \| 79/24222 | response to cyclopentenone |
| H12 | Biological process | GO:0009414 | 2.051e-02 | 8/196 \| 189/24222 | response to water deprivation |
| H12 | Cellular Component | GO:0000151 | 4.475e-02 | 4/182 \| 79/23421 | ubiquitin ligase COmplex |
| iHS | Cellular Component | GO:0009941 | 4.334e-02 | 10/211 \| 409/23421 | chloroplast envelope |
| iHS | Cellular Component | GO:0005737 | 2.780e-02 | 15/211 \| 703/23421 | cytoplasm |
| iHS | Biological process | GO:0042742 | 2.967e-03 | 10/222 \| 176/24222 | defense response to bacterium |
| iHS | Cellular Component | GO:0005663 | 1.257e-02 | 2/211 \| 4/23421 | DNA replication factor C COmplex |
| iHS | Cellular Component | GO:0005615 | 3.254e-02 | 2/211 \| 9/23421 | extracellular space |
| iHS | Cellular Component | GO:0005634 | 1.840e-04 | 42/211 \| 2161/23421 | nucleus |
| iHS | Cellular Component | GO:0009505 | 3.516e-02 | 8/211 \| 262/23421 | plant-type cell wall |
| iHS | Cellular Component | GO:0005886 | 3.645e-03 | 35/211 \| 1929/23421 | plasma membrane |
| iHS | Cellular Component | GO:0043078 | 8.432e-03 | 2/211 \| 3/23421 | polar nucleus |
| iHS | Biological process | GO:0009737 | 7.040e-03 | 12/222 \| 304/24222 | response to aBScisic acid stimulus |
| iHS | Biological process | GO:0009409 | 1.574e-02 | 10/222 \| 243/24222 | response to COld |
| iHS | Biological process | GO:0009826 | 3.895e-02 | 6/222 \| 98/24222 | unidimensional cell growth |
| iHS | Cellular Component | GO:0005773 | 1.699e-02 | 13/211 \| 523/23421 | vacuole |
| LS-17 | Cellular Component | GO:0005618 | 4.422e-02 | 5/49 \| 424/23421 | cell wall |
| LS-17 | Molecular function | GO:0048441 | 4.642e-02 | 2/48 \| 11/24222 | petal development |

Table S9: Hypergeometric test results. Results are rounded to 3 decimals.

| Trait | p-value 99.9th percentile ihs | FDR p-value 99.9th percentile ihs | p-value 99.9th percentile H12 | FDR p-value 99.9th percentile H12 |
|---|---|---|---|---|

| | | | | |
|---|---|---|---|---|
| DBH-17 | 0.200 | 0.338 | 1 | 1 |
| BB2-17 | 0.448 | 0.580 | 0.448 | 0.580 |
| BB4-17 | 1 | 1 | 0.115 | 0.169 |
| CO3-17 | 0.001 | 0.013 | 0 | 0 |
| CO8-17 | 0.078 | 0.231 | 0.747 | 0.865 |
| LS2-17 | 0.003 | 0.031 | 0 | 0 |
| LS5-17 | 1 | 1 | 0 | 0 |
| BB2-18 | 0.327 | 0.450 | 0.085 | 0.134 |
| BB4-18 | 0.099 | 0.231 | 0.076 | 0.134 |
| CO3-18 | 0.042 | 0.186 | 0.042 | 0.112 |
| CO8-18 | 0.055 | 0.203 | 0.046 | 0.112 |
| LS2-18 | 0.269 | 0.394 | 0.269 | 0.370 |
| LS5-18 | 0.017 | 0.095 | 0.002 | 0.011 |
| BB4-brn | 0.140 | 0.275 | 0.036 | 0.112 |
| BB2-stb | 0.105 | 0.231 | 0.085 | 0.134 |
| BB4-stb | 0.017 | 0.095 | 0.010 | 0.037 |
| BB2-stt | 0.150 | 0.275 | 0.876 | 0.963 |
| BB4-stt | 0.475 | 0.581 | 0.476 | 0.581 |
| BB2-top | 0.993 | 1 | 0.064 | 0.134 |
| BB4-top | 0.092 | 0.231 | 0 | 0 |
| BS2 | 1 | 1 | 1 | 1 |

Table S10: All candidate genes identified within 10 kbp of significant slopes.

| Gene | Trait(s) | # of traits | Function in *Arabidopsis* |
|---|---|---|---|
| Potri.001G209400 | BB2-17, BB2-18 | 2 | Lung seven transmembrane receptor family protein |
| Potri.001G209500 | BB2-17, BB2-18 | 2 | Acyl transferase/acyl hydrolase/lysophospholipase superfamily protein |
| Potri.013G094000 | BB2-18 | 1 | S-locus lectin protein kinase family protein |
| Potri.013G094100 | BB2-18 | 1 | AMP-dependent synthetase and ligase family protein |
| Potri.014G188700 | BB2-18 | 1 | zinc ion binding;nucleic acid binding |
| Potri.014G188850 | BB2-18 | 1 | zinc ion binding;nucleic acid binding |
| Potri.017G148900 | BB2-18 | 1 | Tic22-like family protein |
| Potri.017G149000 | BB2-18 | 1 | WRKY DNA-binding protein 27 |
| Potri.017G149100 | BB2-18 | 1 | NADH-ubiquinone oxidoreductase-related |
| Potri.017G149200 | BB2-18 | 1 | |
| Potri.003G009700 | BB2-stb | 1 | Protein of unknown function (DUF803) |
| Potri.003G009800 | BB2-stb | 1 | COx19 family protein (CHCH motif) |
| Potri.003G009900 | BB2-stb | 1 | pumilio 6 |
| Potri.010G055700 | BB2-stb | 1 | Ankyrin repeat family protein |
| Potri.010G055800 | BB2-stb | 1 | lysine histidine transporter 1 |
| Potri.010G055900 | BB2-stb | 1 | DNA polymerase alpha 2 |
| Potri.010G056000 | BB2-stb | 1 | DNA-binding storekeeper protein-related |
| Potri.010G056100 | BB2-stb | 1 | NAD(P)-binding Rossmann-fold superfamily protein |

| | | | |
|---|---|---|---|
| Potri.013G098100 | BB2-stb | 1 | Disease resistance protein (TIR-NBS-LRR class) family |
| Potri.009G093100 | BB2-stt | 1 | CYCLIN A3;4 |
| Potri.009G093200 | BB2-stt | 1 | TIP41-like family protein |
| Potri.009G093300 | BB2-stt | 1 | zinc finger protein 7 |
| Potri.009G093400 | BB2-stt | 1 | |
| Potri.009G093500 | BB2-stt | 1 | 1,2-alpha-L-fuCOsidases |
| Potri.009G094500 | BB2-stt | 1 | Plant basic secretory protein (BSP) family protein |
| Potri.009G094600 | BB2-stt | 1 | Plant basic secretory protein (BSP) family protein |
| Potri.009G094700 | BB2-stt | 1 | Plant basic secretory protein (BSP) family protein |
| Potri.009G094800 | BB2-stt | 1 | S-methyl-5-thioribose kinase |
| Potri.009G094900 | BB2-stt | 1 | lipase 1 |
| Potri.009G095000 | BB2-stt | 1 | |
| Potri.009G095100 | BB2-stt | 1 | UDP-GlyCOsyltransferase superfamily protein |
| Potri.009G095201 | BB2-stt | 1 | UDP-GlyCOsyltransferase superfamily protein |
| Potri.009G095700 | BB2-stt | 1 | Pseudouridine synthase family protein |
| Potri.009G095800 | BB2-stt | 1 | GroES-like zinc-binding alCOhol dehydrogenase family protein |
| Potri.009G098000 | BB2-stt | 1 | PTEN 2 |
| Potri.009G098100 | BB2-stt | 1 | Granulin repeat cysteine protease family protein |
| Potri.009G098200 | BB2-stt | 1 | don-gluCOsyltransferase 1 |
| Potri.009G098300 | BB2-stt | 1 | UDP-gluCOsyl transferase 73B5 |
| Potri.009G098400 | BB2-stt | 1 | UDP-gluCOsyl transferase 73B5 |
| Potri.012G030900 | BB2-stt | 1 | |
| Potri.012G031000 | BB2-stt | 1 | |
| Potri.004G117500 | BB2-top | 1 | PLC-like phosphodiesterases superfamily protein |
| Potri.006G096400 | BB2-top | 1 | Transducin/WD40 repeat-like superfamily protein |
| Potri.006G096500 | BB2-top | 1 | |
| Potri.006G096600 | BB2-top | 1 | Protein of unknown function (DUF740) |
| Potri.008G197000 | BB2-top | 1 | squamosa promoter binding protein-like 7 |
| Potri.008G197100 | BB2-top | 1 | Major facilitator superfamily protein |
| Potri.017G068200 | BB2-top | 1 | |
| Potri.017G068300 | BB2-top | 1 | |
| Potri.014G071900 | BB4-17 | 1 | AAA-type ATPase family protein |
| Potri.014G072000 | BB4-17 | 1 | cytochrome P450, family 704, subfamily A, polypeptide 2 |
| Potri.014G072100 | BB4-17 | 1 | cytochrome P450, family 704, subfamily A, polypeptide 2 |
| Potri.014G072150 | BB4-17 | 1 | cytochrome P450, family 704, subfamily A, polypeptide 2 |
| Potri.016G011500 | BB4-17 | 1 | |
| Potri.016G011601 | BB4-17 | 1 | Leucine-rich repeat transmembrane protein kinase |
| Potri.016G019400 | BB4-17 | 1 | UDP-GlyCOsyltransferase superfamily protein |
| Potri.016G019500 | BB4-17 | 1 | RNI-like superfamily protein |
| Potri.016G019600 | BB4-17 | 1 | receptor like protein 21 |
| Potri.016G019700 | BB4-17 | 1 | Cysteine proteinases superfamily protein |
| Potri.016G026366 | BB4-17 | 1 | PentatriCOpeptide repeat (PPR) superfamily protein |

| | | | |
|---|---|---|---|
| Potri.016G026532 | BB4-17 | 1 | |
| Potri.016G027101 | BB4-17 | 1 | GRAS family transcription factor family protein |
| Potri.016G030800 | BB4-17 | 1 | protein kinase family protein / protein phosphatase 2C ( PP2C) family protein |
| Potri.016G030900 | BB4-17 | 1 | Cystatin/monellin family protein |
| Potri.016G031000 | BB4-17 | 1 | cytochrome P450, family 98, subfamily A, polypeptide 3 |
| Potri.016G137500 | BB4-17 | 1 | VACUOLAR SORTING RECEPTOR 2 |
| Potri.016G137600 | BB4-17 | 1 | cytochrome P450, family 71, subfamily A, polypeptide 25 |
| Potri.016G137700 | BB4-17 | 1 | Lung seven transmembrane receptor family protein |
| Potri.016G137800 | BB4-17 | 1 | Protein of unknown function, DUF599 |
| Potri.016G030500 | BB4-17, BB4-18 | 2 | DNA/RNA helicase protein |
| Potri.016G030600 | BB4-17, BB4-18 | 2 | DNA/RNA helicase protein |
| Potri.016G030700 | BB4-17, BB4-18 | 2 | |
| Potri.016G007101 | BB4-17, BS7 | 2 | |
| Potri.016G007200 | BB4-17, BS7 | 2 | Outward rectifying potassium channel protein |
| Potri.016G007300 | BB4-17, BS7 | 2 | FAR1-related sequence 10 |
| Potri.016G007400 | BB4-17, BS7 | 2 | |
| Potri.016G007500 | BB4-17, BS7 | 2 | agenet domain-COntaining protein / bromo-adjacent homology (BAH) domain-COntaining protein |
| Potri.001G160700 | BB4-18 | 1 | zinc finger (C2H2 type) family protein |
| Potri.001G160800 | BB4-18 | 1 | GTP-binding family protein |
| Potri.002G112700 | BB4-18 | 1 | homolog of yeast ergosterol28 |
| Potri.002G112800 | BB4-18 | 1 | |
| Potri.002G112900 | BB4-18 | 1 | |
| Potri.002G113000 | BB4-18 | 1 | alpha/beta-Hydrolases superfamily protein |
| Potri.015G099200 | BB4-18 | 1 | WRKY DNA-binding protein 75 |
| Potri.015G099400 | BB4-18 | 1 | |
| Potri.015G099500 | BB4-18 | 1 | Heavy metal transport/detoxification superfamily protein |
| Potri.016G030000 | BB4-18 | 1 | Leucine-rich repeat protein kinase family protein |
| Potri.016G030100 | BB4-18 | 1 | syntaxin of plants 51 |
| Potri.016G030200 | BB4-18 | 1 | syntaxin of plants 52 |
| Potri.016G030300 | BB4-18 | 1 | Ubiquitin carboxyl-terminal hydrolase family protein |
| Potri.016G030400 | BB4-18 | 1 | Transducin/WD40 repeat-like superfamily protein |
| Potri.019G106700 | BB4-18 | 1 | Ankyrin repeat family protein |
| Potri.019G106800 | BB4-18 | 1 | |
| Potri.019G106900 | BB4-18 | 1 | |
| Potri.019G106950 | BB4-18 | 1 | |
| Potri.001G275800 | BB4-brn | 1 | Calcium-dependent lipid-binding (CaLB domain) family protein |
| Potri.001G275900 | BB4-brn | 1 | RING/U-box superfamily protein |
| Potri.001G276000 | BB4-brn | 1 | |
| Potri.001G276050 | BB4-brn | 1 | |
| Potri.002G188300 | BB4-brn | 1 | Chaperone DnaJ-domain superfamily protein |
| Potri.002G188500 | BB4-brn | 1 | |

| | | | |
|---|---|---|---|
| Potri.002G188600 | BB4-brn | 1 | Protein kinase superfamily protein |
| Potri.005G143700 | BB4-brn | 1 | Protein kinase superfamily protein |
| Potri.006G132700 | BB4-brn | 1 | TetratriCOpeptide repeat (TPR)-like superfamily protein |
| Potri.006G132800 | BB4-brn | 1 | Zinc finger C-x8-C-x5-C-x3-H type family protein |
| Potri.006G132850 | BB4-brn | 1 | |
| Potri.006G132901 | BB4-brn | 1 | |
| Potri.017G006900 | BB4-brn | 1 | Endonuclease/exonuclease/phosphatase family protein |
| Potri.017G007000 | BB4-brn | 1 | S-adenosyl-L-methionine-dependent methyltransferases superfamily protein |
| Potri.017G007100 | BB4-brn | 1 | |
| Potri.017G007200 | BB4-brn | 1 | Transcription factor IIIC, subunit 5 |
| Potri.017G007300 | BB4-brn | 1 | chloroplast outer envelope protein 37 |
| Potri.017G007400 | BB4-brn | 1 | RNA-binding protein |
| Potri.018G008100 | BB4-brn | 1 | GDSL-like Lipase/Acylhydrolase superfamily protein |
| Potri.018G008200 | BB4-brn | 1 | |
| Potri.018G008300 | BB4-brn | 1 | Protein of unknown function (DUF1262) |
| Potri.018G008400 | BB4-brn | 1 | exocyst subunit exo70 family protein G1 |
| Potri.018G132700 | BB4-brn | 1 | phosphorylethanolamine cytidylyltransferase 1 |
| Potri.018G132800 | BB4-brn | 1 | F-box family protein |
| Potri.018G132900 | BB4-brn | 1 | Leucine-rich repeat transmembrane protein kinase |
| Potri.018G132950 | BB4-brn | 1 | |
| Potri.005G143400 | BB4-brn, BB4-stb | 2 | Protein kinase superfamily protein |
| Potri.005G143600 | BB4-brn, BB4-stb | 2 | Protein kinase superfamily protein |
| Potri.001G205200 | BB4-stb | 1 | Auxin efflux carrier family protein |
| Potri.005G151800 | BB4-stb | 1 | Di-gluCOse binding protein with Kinesin motor domain |
| Potri.005G151900 | BB4-stb | 1 | |
| Potri.005G152000 | BB4-stb | 1 | |
| Potri.005G152100 | BB4-stb | 1 | HVA22 homologue C |
| Potri.005G152200 | BB4-stb | 1 | Sas10/Utp3/C1D family |
| Potri.006G060800 | BB4-stb | 1 | serine-rich protein-related |
| Potri.006G060900 | BB4-stb | 1 | Major facilitator superfamily protein |
| Potri.007G096200 | BB4-stb | 1 | Peroxidase superfamily protein |
| Potri.007G096300 | BB4-stb | 1 | Ribosomal protein S6e |
| Potri.007G096400 | BB4-stb | 1 | Protein kinase superfamily protein |
| Potri.010G016200 | BB4-stb | 1 | Small nuclear RNA activating COmplex (SNAPc), subunit SNAP43 protein |
| Potri.010G016300 | BB4-stb | 1 | Raffinose synthase family protein |
| Potri.010G016400 | BB4-stb | 1 | Raffinose synthase family protein |
| Potri.010G066500 | BB4-stb | 1 | HAD superfamily, subfamily IIIB acid phosphatase |
| Potri.010G066600 | BB4-stb | 1 | seed gene 1 |
| Potri.010G066700 | BB4-stb | 1 | Eukaryotic initiation factor 4E protein |
| Potri.010G066800 | BB4-stb | 1 | |
| Potri.010G066900 | BB4-stb | 1 | proton gradient regulation 5 |

| | | | |
|---|---|---|---|
| Potri.010G067000 | BB4-stb | 1 | Family of unknown function (DUF716) |
| Potri.010G067100 | BB4-stb | 1 | actin-related protein C2B |
| Potri.012G132600 | BB4-stb | 1 | AGAMOUS-like 6 |
| Potri.012G132700 | BB4-stb | 1 | pfkB-like carbohydrate kinase family protein |
| Potri.012G132801 | BB4-stb | 1 | |
| Potri.014G000100 | BB4-stb | 1 | zinc ion binding;nucleic acid binding |
| Potri.014G000200 | BB4-stb | 1 | |
| Potri.019G053500 | BB4-stb | 1 | zinc finger (C2H2 type) family protein |
| Potri.019G053600 | BB4-stb | 1 | ARM repeat superfamily protein |
| Potri.014G047300 | BB4-stt | 1 | Leucine carboxyl methyltransferase |
| Potri.014G047400 | BB4-stt | 1 | Protein of unknown function (DUF630 and DUF632) |
| Potri.018G082100 | BB4-stt | 1 | Protein of unknown function (DUF502) |
| Potri.018G082200 | BB4-stt | 1 | |
| Potri.018G082300 | BB4-stt | 1 | |
| Potri.002G252700 | BB4-top | 1 | HIT-type Zinc finger family protein |
| Potri.002G252800 | BB4-top | 1 | phytochrome interacting factor 3-like 5 |
| Potri.002G252900 | BB4-top | 1 | heat shock protein 60 |
| Potri.002G253000 | BB4-top | 1 | response regulator 24 |
| Potri.002G253100 | BB4-top | 1 | response regulator 24 |
| Potri.002G253200 | BB4-top | 1 | CLIP-associated protein |
| Potri.002G254300 | BB4-top | 1 | myosin XI B |
| Potri.002G254700 | BB4-top | 1 | Transducin family protein / WD-40 repeat family protein |
| Potri.002G254800 | BB4-top | 1 | RmlC-like cupins superfamily protein |
| Potri.002G254900 | BB4-top | 1 | GroES-like zinc-binding dehydrogenase family protein |
| Potri.002G255000 | BB4-top | 1 | ammonium transporter 1;1 |
| Potri.002G255100 | BB4-top | 1 | ammonium transporter 1;1 |
| Potri.002G255200 | BB4-top | 1 | |
| Potri.002G255300 | BB4-top | 1 | TetratriCOpeptide repeat (TPR)-like superfamily protein |
| Potri.002G255400 | BB4-top | 1 | RPM1 interacting protein 13 |
| Potri.002G255900 | BB4-top | 1 | NAD(P)-binding Rossmann-fold superfamily protein |
| Potri.002G256001 | BB4-top | 1 | |
| Potri.002G256100 | BB4-top | 1 | |
| Potri.002G256200 | BB4-top | 1 | Exostosin family protein |
| Potri.002G256300 | BB4-top | 1 | Subtilase family protein |
| Potri.002G256400 | BB4-top | 1 | photosystem II reaction center PSB28 protein |
| Potri.002G256500 | BB4-top | 1 | Leucine-rich repeat transmembrane protein kinase family protein |
| Potri.002G256600 | BB4-top | 1 | indole-3-acetic acid inducible 11 |
| Potri.002G256700 | BB4-top | 1 | |
| Potri.002G258200 | BB4-top | 1 | Leucine-rich repeat receptor-like protein kinase family protein |
| Potri.002G258300 | BB4-top | 1 | |
| Potri.002G262100 | BB4-top | 1 | |
| Potri.002G262200 | BB4-top | 1 | |

| | | | |
|---|---|---|---|
| Potri.002G262300 | BB4-top | 1 | GYF domain-COntaining protein |
| Potri.002G263450 | BB4-top | 1 | |
| Potri.002G263500 | BB4-top | 1 | |
| Potri.004G013100 | BB4-top | 1 | Mitochondrial transcription termination factor family protein |
| Potri.004G013200 | BB4-top | 1 | glyoxalase I homolog |
| Potri.004G013300 | BB4-top | 1 | PentatriCOpeptide repeat (PPR) superfamily protein |
| Potri.004G013400 | BB4-top | 1 | arogenate dehydratase 1 |
| Potri.004G162800 | BB4-top | 1 | ATP binding microtubule motor family protein |
| Potri.004G162966 | BB4-top | 1 | LYR family of Fe/S cluster biogenesis protein |
| Potri.004G163032 | BB4-top | 1 | |
| Potri.004G163100 | BB4-top | 1 | |
| Potri.004G163200 | BB4-top | 1 | ubiquitin fusion degradation 1 |
| Potri.010G035800 | BB4-top | 1 | AAA-type ATPase family protein / ankyrin repeat family protein |
| Potri.010G035900 | BB4-top | 1 | peptidase M20/M25/M40 family protein |
| Potri.013G130400 | BB4-top | 1 | GlyCOsyl hydrolase superfamily protein |
| Potri.013G130500 | BB4-top | 1 | COre-2/I-branching beta-1,6-N-acetylgluCOsaminyltransferase family protein |
| Potri.013G130600 | BB4-top | 1 | PentatriCOpeptide repeat (PPR) superfamily protein |
| Potri.013G130700 | BB4-top | 1 | |
| Potri.017G006400 | BB4-top | 1 | Pectin lyase-like superfamily protein |
| Potri.017G006500 | BB4-top | 1 | Pectin lyase-like superfamily protein |
| Potri.018G041900 | BB4-top | 1 | casein kinase I-like 3 |
| Potri.018G041950 | BB4-top | 1 | casein kinase I-like 3 |
| Potri.018G042000 | BB4-top | 1 | Plant protein of unknown function (DUF828) with plant pleckstrin homology-like region |
| Potri.018G042100 | BB4-top | 1 | Glyoxalase/Bleomycin resistance protein/Dioxygenase superfamily protein |
| Potri.001G100200 | BS2 | 1 | beta-hydroxylase 1 |
| Potri.001G100300 | BS2 | 1 | |
| Potri.001G100400 | BS2 | 1 | |
| Potri.001G100500 | BS2 | 1 | Nuclear transport factor 2 (NTF2) family protein |
| Potri.001G100600 | BS2 | 1 | PentatriCOpeptide repeat (PPR) superfamily protein |
| Potri.001G100700 | BS2 | 1 | Protein of unknown function (DUF604) |
| Potri.002G231400 | BS2 | 1 | |
| Potri.002G231450 | BS2 | 1 | lipase class 3 family protein |
| Potri.002G231501 | BS2 | 1 | lipase class 3 family protein |
| Potri.003G075500 | BS2 | 1 | Pectin lyase-like superfamily protein |
| Potri.003G075600 | BS2 | 1 | PLATZ transcription factor family protein |
| Potri.004G117100 | BS2 | 1 | COBRA-like protein 1 precursor |
| Potri.004G117200 | BS2 | 1 | COBRA-like extracellular glyCOsyl-phosphatidyl inositol-anchored protein family |
| Potri.004G117300 | BS2 | 1 | B12D protein |
| Potri.004G117400 | BS2 | 1 | Heat shock protein 70 (Hsp 70) family protein |
| Potri.006G116000 | BS2 | 1 | demeter-like 1 |

| | | | |
|---|---|---|---|
| Potri.006G116200 | BS2 | 1 | Ergosterol biosynthesis ERG4/ERG24 family |
| Potri.006G116300 | BS2 | 1 | |
| Potri.006G117900 | BS2 | 1 | Sulfite exporter TauE/SafE family protein |
| Potri.006G118000 | BS2 | 1 | Exostosin family protein |
| Potri.006G118100 | BS2 | 1 | ENTH/ANTH/VHS superfamily protein |
| Potri.014G071100 | BS2 | 1 | dihydrodipiCOlinate synthase |
| Potri.014G071200 | BS2 | 1 | protein binding |
| Potri.014G071300 | BS2 | 1 | |
| Potri.014G071400 | BS2 | 1 | SMAD/FHA domain-COntaining protein |
| Potri.014G071500 | BS2 | 1 | NB-ARC domain-COntaining disease resistance protein |
| Potri.017G023200 | BS2 | 1 | Ypt/Rab-GAP domain of gyp1p superfamily protein |
| Potri.017G023300 | BS2 | 1 | Protein of unknown function (DUF3755) |
| Potri.017G039700 | BS2 | 1 | cysteine-rich RLK (RECEPTOR-like protein kinase) 16 |
| Potri.017G039800 | BS2 | 1 | AGAMOUS-like 62 |
| Potri.017G039900 | BS2 | 1 | Receptor-like protein kinase-related family protein |
| Potri.017G040000 | BS2 | 1 | Receptor-like protein kinase-related family protein |
| Potri.019G071900 | BS2 | 1 | SNARE associated Golgi protein family |
| Potri.019G072100 | BS2 | 1 | |
| Potri.019G072200 | BS2 | 1 | |
| Potri.019G072300 | BS2 | 1 | shoot gravitropism 2 (SGR2) |
| Potri.019G072400 | BS2 | 1 | |
| Potri.005G021400 | BS7 | 1 | eukaryotic translation initiation factor 3A |
| Potri.005G021500 | BS7 | 1 | |
| Potri.005G021601 | BS7 | 1 | PRP38 family protein |
| Potri.005G021700 | BS7 | 1 | TOPLESS-related 3 |
| Potri.005G026400 | BS7 | 1 | RECQ helicase SIM |
| Potri.005G026450 | BS7 | 1 | |
| Potri.005G026500 | BS7 | 1 | histone H2A 7 |
| Potri.005G026600 | BS7 | 1 | Transducin/WD40 repeat-like superfamily protein |
| Potri.005G026700 | BS7 | 1 | soybean gene regulated by COld-2 |
| Potri.013G000800 | BS7 | 1 | Major facilitator superfamily protein |
| Potri.013G000900 | BS7 | 1 | Pectinacetylesterase family protein |
| Potri.013G001000 | BS7 | 1 | myb domain protein 61 |
| Potri.013G001150 | BS7 | 1 | neurofilament protein-related |
| Potri.013G001300 | BS7 | 1 | phytochrome interacting factor 3 |
| Potri.013G003000 | BS7 | 1 | putative endonuclease or glyCOsyl hydrolase with C2H2-type zinc finger domain |
| Potri.013G003100 | BS7 | 1 | Putative endonuclease or glyCOsyl hydrolase |
| Potri.013G003200 | BS7 | 1 | Putative endonuclease or glyCOsyl hydrolase |
| Potri.016G006900 | BS7 | 1 | microtubule-associated proteins 70-5 |
| Potri.016G007000 | BS7 | 1 | |
| Potri.017G145901 | BS7 | 1 | Pectin lyase-like superfamily protein |

| | | | |
|---|---|---|---|
| Potri.017G146000 | BS7 | 1 | Protein kinase superfamily protein |
| Potri.017G146100 | BS7 | 1 | |
| Potri.017G146200 | BS7 | 1 | glutaminyl cyclase |
| Potri.017G146300 | BS7 | 1 | F-box/RNI-like superfamily protein |
| Potri.017G146400 | BS7 | 1 | F-box/RNI-like superfamily protein |
| Potri.010G043100 | CO3-17 | 1 | |
| Potri.010G043200 | CO3-17 | 1 | NSP-interacting kinase 3 |
| Potri.010G043300 | CO3-17 | 1 | Plant protein of unknown function (DUF828) |
| Potri.010G043400 | CO3-17 | 1 | |
| Potri.010G162200 | CO3-17 | 1 | GlyCOsyltransferase family 61 protein |
| Potri.010G162300 | CO3-17 | 1 | chorismate mutase 3 |
| Potri.010G162400 | CO3-17 | 1 | Protein of unknown function, DUF538 |
| Potri.010G162500 | CO3-17 | 1 | Protein of unknown function (DUF155) |
| Potri.010G162600 | CO3-17 | 1 | homologue of bacterial MinE 1 |
| Potri.010G162700 | CO3-17 | 1 | Transducin/WD40 repeat-like superfamily protein |
| Potri.010G167100 | CO3-17 | 1 | RNA-binding KH domain-COntaining protein |
| Potri.010G167150 | CO3-17 | 1 | |
| Potri.010G167200 | CO3-17 | 1 | expansin A1 |
| Potri.010G172900 | CO3-17 | 1 | Cation efflux family protein |
| Potri.010G173000 | CO3-17 | 1 | Uncharacterized COnserved protein (DUF2358) |
| Potri.010G173100 | CO3-17 | 1 | Protein kinase superfamily protein |
| Potri.010G180000 | CO3-17 | 1 | HXXXD-type acyl-transferase family protein |
| Potri.010G180100 | CO3-17 | 1 | |
| Potri.010G180200 | CO3-17 | 1 | |
| Potri.010G180300 | CO3-17 | 1 | Cysteine proteinases superfamily protein |
| Potri.010G180400 | CO3-17 | 1 | 2-phosphoglyCOlate phosphatase 1 |
| Potri.010G180500 | CO3-17 | 1 | UDP-galactose transporter 6 |
| Potri.010G184500 | CO3-17 | 1 | ferritin 2 |
| Potri.010G184600 | CO3-17 | 1 | histidine acid phosphatase family protein |
| Potri.010G184700 | CO3-17 | 1 | PLAC8 family protein |
| Potri.010G184800 | CO3-17 | 1 | XH/XS domain-COntaining protein |
| Potri.010G184900 | CO3-17 | 1 | Eukaryotic translation initiation factor 2 subunit 1 |
| Potri.010G185000 | CO3-17 | 1 | S-adenosyl-L-methionine-dependent methyltransferases superfamily protein |
| Potri.010G185100 | CO3-17 | 1 | Peptide-N4-(N-acetyl-beta-gluCOsaminyl)asparagine amidase A protein |
| Potri.010G185200 | CO3-17 | 1 | Ankyrin repeat family protein |
| Potri.010G205500 | CO3-17 | 1 | heat shock COgnate protein 70-1 |
| Potri.010G205600 | CO3-17 | 1 | Heat shock protein 70 (Hsp 70) family protein |
| Potri.010G205700 | CO3-17 | 1 | heat shock protein 70 |
| Potri.012G070600 | CO3-17 | 1 | Protein of unknown function (DUF1118) |
| Potri.012G070700 | CO3-17 | 1 | nitrate transporter 1:2 |
| Potri.012G070801 | CO3-17 | 1 | |

| | | | |
|---|---|---|---|
| Potri.013G118500 | CO3-17 | 1 | TRICHOME BIREFRINGENCE-LIKE 39 |
| Potri.013G118600 | CO3-17 | 1 | late embryogenesis abundant domain-COntaining protein / LEA domain-COntaining protein |
| Potri.013G118700 | CO3-17 | 1 | UDP-gluCOsyl transferase 78D2 |
| Potri.013G118800 | CO3-17 | 1 | |
| Potri.013G118900 | CO3-17 | 1 | global transcription factor group E4 |
| Potri.013G147600 | CO3-17 | 1 | zinc ion binding |
| Potri.013G147700 | CO3-17 | 1 | Wound-responsive family protein |
| Potri.013G147750 | CO3-17 | 1 | |
| Potri.013G147800 | CO3-17 | 1 | PapD-like superfamily protein |
| Potri.013G147900 | CO3-17 | 1 | Wound-responsive family protein |
| Potri.013G148000 | CO3-17 | 1 | Wound-responsive family protein |
| Potri.004G228800 | CO3-17, CO3-18 | 2 | auxin response factor 1 |
| Potri.004G228900 | CO3-17, CO3-18 | 2 | mitotic checkpoint family protein |
| Potri.002G013700 | CO3-17, CO3-18, LS2-17, LS5-18 | 4 | SKU5 similar 5 |
| Potri.002G013800 | CO3-17, CO3-18, LS2-17, LS5-18 | 4 | enoyl-COA hydratase 2 |
| Potri.002G013900 | CO3-17, CO3-18, LS2-17, LS5-18 | 4 | Prolyl oligopeptidase family protein |
| Potri.002G014000 | CO3-17, CO3-18, LS2-17, LS5-18 | 4 | Prolyl oligopeptidase family protein |
| Potri.002G014100 | CO3-17, CO3-18, LS2-17, LS5-18 | 4 | rho guanyl-nucleotide exchange factor 1 |
| Potri.010G185800 | CO3-17, CO8-17 | 2 | UDP-gluCOse pyrophosphorylase 3 |
| Potri.010G185900 | CO3-17, CO8-17 | 2 | plant U-box 13 |
| Potri.002G014200 | CO3-17, LS5-18 | 2 | |
| Potri.003G044800 | CO3-18 | 1 | |
| Potri.003G044900 | CO3-18 | 1 | AAA-type ATPase family protein |
| Potri.016G043500 | CO3-18 | 1 | INO80 ortholog |
| Potri.016G043600 | CO3-18 | 1 | Calcium-binding EF-hand family protein |
| Potri.016G043700 | CO3-18 | 1 | alpha carbonic anhydrase 4 |
| Potri.016G043800 | CO3-18 | 1 | eukaryotic translation initiation factor 3E |
| Potri.002G037900 | CO3-18, LS5-18 | 2 | Peptidase S24/S26A/S26B/S26C family protein |
| Potri.002G038000 | CO3-18, LS5-18 | 2 | ENTH/VHS/GAT family protein |
| Potri.002G038100 | CO3-18, LS5-18 | 2 | RmlC-like cupins superfamily protein |
| Potri.002G038200 | CO3-18, LS5-18 | 2 | |
| Potri.002G038300 | CO3-18, LS5-18 | 2 | translocase inner membrane subunit 44-2 |
| Potri.003G159500 | CO3-18, LS5-18 | 2 | Translation protein SH3-like family protein |
| Potri.003G159600 | CO3-18, LS5-18 | 2 | Protein phosphatase 2C family protein |
| Potri.003G159700 | CO3-18, LS5-18 | 2 | xyloglucan endotransgluCOsylase/hydrolase 5 |
| Potri.003G159750 | CO3-18, LS5-18 | 2 | |
| Potri.007G077300 | CO3-18, LS5-18 | 2 | |
| Potri.007G077400 | CO3-18, LS5-18 | 2 | essential meiotic endonuclease 1B |
| Potri.007G077500 | CO3-18, LS5-18 | 2 | Protein of unknown function (DUF3511) |

| | | | |
|---|---|---|---|
| Potri.007G077600 | CO3-18, LS5-18 | 2 | Protein kinase superfamily protein |
| Potri.007G077700 | CO3-18, LS5-18 | 2 | |
| Potri.009G162601 | CO8-17 | 1 | nitrilase-like protein 1 |
| Potri.009G162700 | CO8-17 | 1 | |
| Potri.009G162800 | CO8-17 | 1 | Protein of unknown function (DUF3049) |
| Potri.009G162900 | CO8-17 | 1 | Rho GTPase activating protein with PAK-box/P21-Rho-binding domain |
| Potri.009G163001 | CO8-17 | 1 | |
| Potri.009G163100 | CO8-17 | 1 | COpper ion binding |
| Potri.009G163200 | CO8-17 | 1 | COpper/zinc superoxide dismutase 2 |
| Potri.017G092100 | CO8-17 | 1 | TPX2 (targeting protein for Xklp2) protein family |
| Potri.017G092200 | CO8-17 | 1 | Ribosomal protein S19e family protein |
| Potri.017G093600 | CO8-17 | 1 | RS2-interacting KH protein |
| Potri.017G093650 | CO8-17 | 1 | exocyst subunit exo70 family protein E1 |
| Potri.017G093700 | CO8-17 | 1 | exocyst subunit exo70 family protein E1 |
| Potri.017G093800 | CO8-17 | 1 | exocyst subunit exo70 family protein E1 |
| Potri.019G061800 | CO8-17 | 1 | serine/threonine protein kinase 2 |
| Potri.019G061900 | CO8-17 | 1 | phosphate transporter 1;3 |
| Potri.019G062000 | CO8-17 | 1 | thioredoxin H-type 9 |
| Potri.019G062100 | CO8-17 | 1 | Leucine-rich repeat protein kinase family protein |
| Potri.019G062200 | CO8-17 | 1 | |
| Potri.013G025900 | CO8-17, LS5-17 | 2 | basic helix-loop-helix (bHLH) DNA-binding superfamily protein |
| Potri.013G026000 | CO8-17, LS5-17 | 2 | NAD(P)-binding Rossmann-fold superfamily protein |
| Potri.013G026100 | CO8-17, LS5-17 | 2 | NAD(P)-binding Rossmann-fold superfamily protein |
| Potri.001G137800 | CO8-18 | 1 | homeobox-7 |
| Potri.002G203300 | CO8-18 | 1 | Protein of unknown function, DUF538 |
| Potri.002G203400 | CO8-18 | 1 | Protein of unknown function, DUF538 |
| Potri.002G230400 | CO8-18 | 1 | Late embryogenesis abundant (LEA) hydroxyproline-rich glyCOprotein family |
| Potri.002G230500 | CO8-18 | 1 | DNA mismatch repair protein MutS, type 2 |
| Potri.002G230600 | CO8-18 | 1 | Eukaryotic protein of unknown function (DUF914) |
| Potri.019G129600 | CO8-18 | 1 | |
| Potri.019G129700 | CO8-18 | 1 | |
| Potri.019G129800 | CO8-18 | 1 | ChaC-like family protein |
| Potri.019G129900 | CO8-18 | 1 | chromatin remodeling 1 |
| Potri.019G130000 | CO8-18 | 1 | Basic-leucine zipper (bZIP) transcription factor family protein |
| Potri.009G106300 | CO8-18, LS5-18 | 2 | |
| Potri.009G106400 | CO8-18, LS5-18 | 2 | Peroxidase superfamily protein |
| Potri.009G106500 | CO8-18, LS5-18 | 2 | RNA-binding protein |
| Potri.001G392901 | DBH17 | 1 | UDP-GlyCOsyltransferase superfamily protein |
| Potri.001G393001 | DBH17 | 1 | |
| Potri.001G393101 | DBH17 | 1 | |
| Potri.006G031700 | DBH17 | 1 | disproportionating enzyme 2 |

| | | | |
|---|---|---|---|
| Potri.006G031800 | DBH17 | 1 | hydroxyproline-rich glyCOprotein family protein |
| Potri.006G031900 | DBH17 | 1 | global transcription factor group B1 |
| Potri.011G143700 | DBH17 | 1 | GTP-binding protein-related |
| Potri.011G143800 | DBH17 | 1 | methyl esterase 14 |
| Potri.011G143900 | DBH17 | 1 | |
| Potri.001G368300 | LS2-17 | 1 | COncanavalin A-like lectin protein kinase family protein |
| Potri.001G368400 | LS2-17 | 1 | Polynucleotidyl transferase, ribonuclease H-like superfamily protein |
| Potri.001G368500 | LS2-17 | 1 | NOL1/NOP2/sun family protein / antitermination NusB domain-COntaining protein |
| Potri.001G373200 | LS2-17 | 1 | GroES-like zinc-binding alCOhol dehydrogenase family protein |
| Potri.001G373300 | LS2-17 | 1 | sequence-specific DNA binding transcription factors;sequence-specific DNA binding |
| Potri.001G373400 | LS2-17 | 1 | |
| Potri.001G374500 | LS2-17 | 1 | alpha/beta-Hydrolases superfamily protein |
| Potri.001G374600 | LS2-17 | 1 | glutamate receptor 2.7 |
| Potri.001G375300 | LS2-17 | 1 | sodium/calcium exchanger family protein / calcium-binding EF hand family protein |
| Potri.001G375400 | LS2-17 | 1 | sodium/calcium exchanger family protein / calcium-binding EF hand family protein |
| Potri.001G375700 | LS2-17 | 1 | IQ-domain 10 |
| Potri.001G375800 | LS2-17 | 1 | TCP family transcription factor 4 |
| Potri.001G375900 | LS2-17 | 1 | P-loop COntaining nucleoside triphosphate hydrolases superfamily protein |
| Potri.001G376900 | LS2-17 | 1 | |
| Potri.001G377000 | LS2-17 | 1 | Major facilitator superfamily protein |
| Potri.001G377100 | LS2-17 | 1 | |
| Potri.001G377200 | LS2-17 | 1 | DNA-directed RNA polymerase II |
| Potri.001G377300 | LS2-17 | 1 | |
| Potri.001G377400 | LS2-17 | 1 | RNA-binding (RRM/RBD/RNP motifs) family protein |
| Potri.001G378000 | LS2-17 | 1 | RING/U-box superfamily protein |
| Potri.003G103100 | LS2-17 | 1 | |
| Potri.003G103200 | LS2-17 | 1 | TRAF-like family protein |
| Potri.003G103300 | LS2-17 | 1 | CLP protease P4 |
| Potri.003G103400 | LS2-17 | 1 | Ankyrin repeat family protein |
| Potri.003G103500 | LS2-17 | 1 | NAC-like, activated by AP3/PI |
| Potri.003G103600 | LS2-17 | 1 | Amino acid permease family protein |
| Potri.003G111700 | LS2-17 | 1 | Phosphatidic acid phosphatase (PAP2) family protein |
| Potri.006G214300 | LS2-17 | 1 | exocyst subunit exo70 family protein H4 |
| Potri.006G214400 | LS2-17 | 1 | Pectin lyase-like superfamily protein |
| Potri.006G214500 | LS2-17 | 1 | Ankyrin repeat family protein |
| Potri.007G101100 | LS2-17 | 1 | ssDNA-binding transcriptional regulator |
| Potri.007G101201 | LS2-17 | 1 | wound-responsive family protein |
| Potri.007G101300 | LS2-17 | 1 | |
| Potri.009G021400 | LS2-17 | 1 | HSP20-like chaperones superfamily protein |

| | | | |
|---|---|---|---|
| Potri.009G021500 | LS2-17 | 1 | calmodulin 5 |
| Potri.009G021600 | LS2-17 | 1 | 5\'-AMP-activated protein kinase beta-2 subunit protein |
| Potri.013G100600 | LS2-17 | 1 | Nuclear transport factor 2 (NTF2) family protein |
| Potri.016G033300 | LS2-17 | 1 | Transcription elongation factor (TFIIS) family protein |
| Potri.016G033800 | LS2-17 | 1 | DNA/RNA helicase protein |
| Potri.016G033900 | LS2-17 | 1 | DNA/RNA helicase protein |
| Potri.016G034000 | LS2-17 | 1 | DNA/RNA helicase protein |
| Potri.016G034100 | LS2-17 | 1 | IQ calmodulin-binding motif family protein |
| Potri.016G034200 | LS2-17 | 1 | AtGCP3 interacting protein 1 |
| Potri.016G034300 | LS2-17 | 1 | serine carboxypeptidase-like 45 |
| Potri.016G033400 | LS2-17, LS2-18 | 2 | |
| Potri.016G033500 | LS2-17, LS2-18 | 2 | DNA/RNA helicase protein |
| Potri.016G033600 | LS2-17, LS2-18 | 2 | DNA/RNA helicase protein |
| Potri.016G033700 | LS2-17, LS2-18 | 2 | DNA/RNA helicase protein |
| Potri.003G213800 | LS2-18 | 1 | fatty acid desaturase 2 |
| Potri.003G213900 | LS2-18 | 1 | |
| Potri.003G214000 | LS2-18 | 1 | C2H2 and C2HC zinc fingers superfamily protein |
| Potri.003G214100 | LS2-18 | 1 | Transducin/WD40 repeat-like superfamily protein |
| Potri.003G214200 | LS2-18 | 1 | glucan synthase-like 12 |
| Potri.006G232700 | LS2-18 | 1 | Protein phosphatase 2C family protein |
| Potri.006G232800 | LS2-18 | 1 | |
| Potri.006G232900 | LS2-18 | 1 | gamma vacuolar processing enzyme |
| Potri.006G233000 | LS2-18 | 1 | |
| Potri.010G111900 | LS2-18 | 1 | P-loop COntaining nucleoside triphosphate hydrolases superfamily protein |
| Potri.010G112000 | LS2-18 | 1 | nudix hydrolase homolog 25 |
| Potri.010G112050 | LS2-18 | 1 | |
| Potri.001G034100 | LS5-17 | 1 | fuCOsyltransferase 1 |
| Potri.001G034200 | LS5-17 | 1 | Ubiquitin-COnjugating enzyme/RWD-like protein |
| Potri.001G034300 | LS5-17 | 1 | global transcription factor group B1 |
| Potri.001G155700 | LS5-17 | 1 | Integrase-type DNA-binding superfamily protein |
| Potri.001G155750 | LS5-17 | 1 | |
| Potri.001G205700 | LS5-17 | 1 | CD2-binding protein-related |
| Potri.001G253000 | LS5-17 | 1 | SELT-like protein precursor |
| Potri.001G253100 | LS5-17 | 1 | Protein of unknown function (DUF803) |
| Potri.001G253200 | LS5-17 | 1 | 3\'-5\'-exoribonuclease family protein |
| Potri.001G253700 | LS5-17 | 1 | phytochromobilin:ferredoxin oxidoreductase, chloroplast / phytochromobilin synthase (HY2) |
| Potri.001G253800 | LS5-17 | 1 | sugar transporter 14 |
| Potri.001G253900 | LS5-17 | 1 | NAD(P)-binding Rossmann-fold superfamily protein |
| Potri.001G254001 | LS5-17 | 1 | indigoidine synthase A family protein |
| Potri.001G255200 | LS5-17 | 1 | |
| Potri.001G255300 | LS5-17 | 1 | |

| | | | |
|---|---|---|---|
| Potri.001G255400 | LS5-17 | 1 | RNA-binding (RRM/RBD/RNP motifs) family protein |
| Potri.001G255466 | LS5-17 | 1 | |
| Potri.001G255532 | LS5-17 | 1 | Zinc finger (CCCH-type) family protein / RNA reCOgnition motif (RRM)-COntaining protein |
| Potri.001G323500 | LS5-17 | 1 | B-box type zinc finger protein with CCT domain |
| Potri.001G323600 | LS5-17 | 1 | Endosomal targeting BRO1-like domain-COntaining protein |
| Potri.001G323700 | LS5-17 | 1 | Uncharacterised protein family (UPF0497) |
| Potri.001G323800 | LS5-17 | 1 | Esterase/lipase/thioesterase family protein |
| Potri.001G323900 | LS5-17 | 1 | |
| Potri.001G324000 | LS5-17 | 1 | Esterase/lipase/thioesterase family protein |
| Potri.001G324100 | LS5-17 | 1 | LRR and NB-ARC domains-COntaining disease resistance protein |
| Potri.001G324200 | LS5-17 | 1 | Esterase/lipase/thioesterase family protein |
| Potri.001G369100 | LS5-17 | 1 | cellulose synthase like E1 |
| Potri.001G369200 | LS5-17 | 1 | Plant protein of unknown function (DUF827) |
| Potri.001G387600 | LS5-17 | 1 | SecY protein transport family protein |
| Potri.002G263800 | LS5-17 | 1 | cytochrome P450, family 721, subfamily A, polypeptide 1 |
| Potri.002G263900 | LS5-17 | 1 | Plant self-inCOmpatibility protein S1 family |
| Potri.002G264000 | LS5-17 | 1 | |
| Potri.003G087600 | LS5-17 | 1 | S-adenosyl-L-methionine-dependent methyltransferases superfamily protein |
| Potri.005G085600 | LS5-17 | 1 | Domain of unknown function (DUF1726) ;Putative ATPase (DUF699) |
| Potri.005G085700 | LS5-17 | 1 | TetratriCOpeptide repeat (TPR)-like superfamily protein |
| Potri.005G085800 | LS5-17 | 1 | violaxanthin de-epoxidase-related |
| Potri.005G085900 | LS5-17 | 1 | S-adenosylmethionine carrier 1 |
| Potri.005G095900 | LS5-17 | 1 | potassium transporter 1 |
| Potri.005G096000 | LS5-17 | 1 | |
| Potri.005G096100 | LS5-17 | 1 | |
| Potri.005G096500 | LS5-17 | 1 | |
| Potri.005G151000 | LS5-17 | 1 | |
| Potri.005G151100 | LS5-17 | 1 | Zinc knuckle (CCHC-type) family protein |
| Potri.006G200150 | LS5-17 | 1 | |
| Potri.006G200200 | LS5-17 | 1 | RING-H2 group F2A |
| Potri.007G018200 | LS5-17 | 1 | |
| Potri.007G018300 | LS5-17 | 1 | |
| Potri.007G018350 | LS5-17 | 1 | |
| Potri.007G018400 | LS5-17 | 1 | Cytochrome P450 superfamily protein |
| Potri.007G045100 | LS5-17 | 1 | Subtilisin-like serine endopeptidase family protein |
| Potri.010G180800 | LS5-17 | 1 | Calcineurin-like metallo-phosphoesterase superfamily protein |
| Potri.010G180900 | LS5-17 | 1 | GDSL-like Lipase/Acylhydrolase family protein |
| Potri.010G181000 | LS5-17 | 1 | Integrase-type DNA-binding superfamily protein |
| Potri.012G034300 | LS5-17 | 1 | little nuclei4 |
| Potri.012G034400 | LS5-17 | 1 | |

| | | | |
|---|---|---|---|
| Potri.012G034500 | LS5-17 | 1 | |
| Potri.012G034600 | LS5-17 | 1 | disease resistance family protein / LRR family protein |
| Potri.012G034700 | LS5-17 | 1 | FAD-binding Berberine family protein |
| Potri.012G034800 | LS5-17 | 1 | Nucleoporin interacting Component (Nup93/Nic96-like) family protein |
| Potri.012G035200 | LS5-17 | 1 | |
| Potri.012G035300 | LS5-17 | 1 | SLAC1 homologue 3 |
| Potri.012G077400 | LS5-17 | 1 | Xanthine/uracil permease family protein |
| Potri.012G105000 | LS5-17 | 1 | Polynucleotide adenylyltransferase family protein |
| Potri.012G105100 | LS5-17 | 1 | phosphate transporter 3;1 |
| Potri.012G105200 | LS5-17 | 1 | AAA-type ATPase family protein |
| Potri.012G105300 | LS5-17 | 1 | acyl carrier protein 4 |
| Potri.013G033000 | LS5-17 | 1 | Esterase/lipase/thioesterase family protein |
| Potri.013G033101 | LS5-17 | 1 | Esterase/lipase/thioesterase family protein |
| Potri.013G033200 | LS5-17 | 1 | Matrixin family protein |
| Potri.013G044100 | LS5-17 | 1 | Signal transduction histidine kinase, hybrid-type, ethylene sensor |
| Potri.013G044200 | LS5-17 | 1 | |
| Potri.013G044300 | LS5-17 | 1 | Methyltransferase family protein |
| Potri.013G044400 | LS5-17 | 1 | |
| Potri.013G060300 | LS5-17 | 1 | calmodulin-binding protein |
| Potri.013G090400 | LS5-17 | 1 | WRKY DNA-binding protein 55 |
| Potri.013G090500 | LS5-17 | 1 | Protein kinase family protein with ARM repeat domain |
| Potri.013G090800 | LS5-17 | 1 | SNF1 kinase homolog 10 |
| Potri.013G098700 | LS5-17 | 1 | Protein kinase family protein with ARM repeat domain |
| Potri.015G009566 | LS5-17 | 1 | ARABIDOPSIS TRITHORAX-RELATED PROTEIN 6 |
| Potri.015G009632 | LS5-17 | 1 | ARABIDOPSIS TRITHORAX-RELATED PROTEIN 6 |
| Potri.015G009700 | LS5-17 | 1 | Transducin/WD40 repeat-like superfamily protein |
| Potri.015G048900 | LS5-17 | 1 | myosin heavy chain-related |
| Potri.015G049000 | LS5-17 | 1 | soluble N-ethylmaleimide-sensitive factor adaptor protein 33 |
| Potri.015G049100 | LS5-17 | 1 | glucuronidase 2 |
| Potri.015G050200 | LS5-17 | 1 | NAD(P)-binding Rossmann-fold superfamily protein |
| Potri.015G050300 | LS5-17 | 1 | Transmembrane Fragile-X-F-associated protein |
| Potri.015G053100 | LS5-17 | 1 | actin-related protein 4 |
| Potri.015G053300 | LS5-17 | 1 | Eukaryotic aspartyl protease family protein |
| Potri.015G053400 | LS5-17 | 1 | aluminum-activated malate transporter 9 |
| Potri.016G098100 | LS5-17 | 1 | C2H2-type zinc finger family protein |
| Potri.019G063400 | LS5-17 | 1 | FAD-binding Berberine family protein |
| Potri.019G087700 | LS5-17 | 1 | somatic embryogenesis receptor-like kinase 1 |
| Potri.019G087800 | LS5-17 | 1 | |
| Potri.019G087900 | LS5-17 | 1 | SU(VAR)3-9 homolog 1 |
| Potri.019G088000 | LS5-17 | 1 | Integrase-type DNA-binding superfamily protein |
| Potri.019G088100 | LS5-17 | 1 | SEC6 |

| | | | |
|---|---|---|---|
| Potri.002G263600 | LS5-17, BB4-top | 2 | indeterminate(ID)-domain 2 |
| Potri.002G263700 | LS5-17, BB4-top | 2 | |
| Potri.003G123600 | LS5-18 | 1 | P-loop COntaining nucleoside triphosphate hydrolases superfamily protein |
| Potri.003G123700 | LS5-18 | 1 | UDP-D-gluCOse/UDP-D-galactose 4-epimerase 1 |
| Potri.003G123800 | LS5-18 | 1 | Homeodomain-like protein |
| Potri.003G123900 | LS5-18 | 1 | Transcription initiation factor TFIIE, beta subunit |
| Potri.003G131500 | LS5-18 | 1 | Nuclear transport factor 2 (NTF2) family protein |
| Potri.003G131550 | LS5-18 | 1 | |
| Potri.003G141200 | LS5-18 | 1 | Glycine-rich protein family |
| Potri.003G141300 | LS5-18 | 1 | SGNH hydrolase-type esterase superfamily protein |
| Potri.003G141400 | LS5-18 | 1 | Calcium-binding EF-hand family protein |
| Potri.003G141500 | LS5-18 | 1 | Galactose-binding protein |
| Potri.003G141600 | LS5-18 | 1 | Nuclear transport factor 2 (NTF2) family protein |
| Potri.003G141700 | LS5-18 | 1 | |
| Potri.007G076700 | LS5-18 | 1 | ribonucleotide reductase 1 |
| Potri.007G076900 | LS5-18 | 1 | 26S proteasome regulatory subunit, putative (RPN5) |
| Potri.009G049700 | LS5-18 | 1 | THUMP domain-COntaining protein |
| Potri.009G049800 | LS5-18 | 1 | HSP20-like chaperones superfamily protein |
| Potri.009G049900 | LS5-18 | 1 | HSP20-like chaperones superfamily protein |
| Potri.009G050000 | LS5-18 | 1 | hexokinase-like 1 |
| Potri.009G092200 | LS5-18 | 1 | SWIB/MDM2 domain superfamily protein |
| Potri.009G092300 | LS5-18 | 1 | arabinogalactan protein 18 |
| Potri.009G092400 | LS5-18 | 1 | Protein of unknown function (DUF620) |
| Potri.009G106200 | LS5-18 | 1 | peptidyl-prolyl cis-trans isomerase / cyclophilin-40 (CYP40) / rotamase |

Table S11: Shared candidate genes between our results and Evans et al. (2014) and McKown et al. (2014).

| Gene | Trait in our results | Source | Trait in source | Year | Description of function in Arabidopis |
|---|---|---|---|---|---|
| Potri.001G137800 | CO8-18 | Evans et al. (2014) | budset | NA | homeobox-7 |
| Potri.001G155700 | LS5-17 | Evans et al. (2014) | bud burst | NA | Integrase-type DNA-binding superfamily protein |
| Potri.001G155700 | LS5-17 | Evans et al. (2014) | height | NA | Integrase-type DNA-binding superfamily protein |
| Potri.001G205200 | BB4-stb | Evans et al. (2014) | budset | NA | Auxin efflux carrier family protein |
| Potri.001G253800 | LS5-17 | Evans et al. (2014) | height | NA | sugar transporter 14 |
| Potri.001G255400 | LS5-17 | Evans et al. (2014) | budset | NA | RNA-binding (RRM/RBD/RNP motifs) family protein |
| Potri.001G275800 | BB4-brn | Evans et al. (2014) | budset | NA | Calcium-dependent lipid-binding (CaLB domain) family protein |
| Potri.001G275900 | BB4-brn | Evans et al. (2014) | budset | NA | RING/U-box superfamily protein |
| Potri.001G323800 | LS5-17 | Evans et al. (2014) | bud burst | NA | Esterase/lipase/thioesterase family protein |

| | | | | | |
|---|---|---|---|---|---|
| Potri.001G323800 | LS5-17 | Evans et al. (2014) | height | NA | Esterase/lipase/thioesterase family protein |
| Potri.001G324000 | LS5-17 | Evans et al. (2014) | bud burst | NA | Esterase/lipase/thioesterase family protein |
| Potri.001G369100 | LS5-17 | Evans et al. (2014) | bud burst | NA | cellulose synthase like E1 |
| Potri.001G369100 | LS5-17 | Evans et al. (2014) | height | NA | cellulose synthase like E1 |
| Potri.001G373300 | LS2-17 | Evans et al. (2014) | budset | NA | sequence-specific DNA binding transcription factors;sequence-specific DNA binding |
| Potri.001G373400 | LS2-17 | Evans et al. (2014) | budset | NA | |
| Potri.001G375400 | LS2-17 | Evans et al. (2014) | budset | NA | sodium/calcium exchanger family protein / calcium-binding EF hand family protein |
| Potri.001G375800 | LS2-17 | Evans et al. (2014) | bud burst | NA | TCP family transcription factor 4 |
| Potri.001G375900 | LS2-17 | Evans et al. (2014) | bud burst | NA | P-loop COntaining nucleoside triphosphate hydrolases superfamily protein |
| Potri.001G377100 | LS2-17 | Evans et al. (2014) | bud burst | NA | |
| Potri.001G377200 | LS2-17 | Evans et al. (2014) | bud burst | NA | DNA-directed RNA polymerase II |
| Potri.001G387600 | LS5-17 | Evans et al. (2014) | budset | NA | SecY protein transport family protein |
| Potri.002G038100 | CO3-18, LS5-18 | Evans et al. (2014) | bud burst | NA | RmlC-like cupins superfamily protein |
| Potri.002G038100 | CO3-18, LS5-18 | Evans et al. (2014) | height | NA | RmlC-like cupins superfamily protein |
| Potri.002G112700 | BB4-18 | Evans et al. (2014) | height | NA | homolog of yeast ergosterol28 |
| Potri.002G113000 | BB4-18 | Evans et al. (2014) | budset | NA | alpha/beta-Hydrolases superfamily protein |
| Potri.002G230600 | CO8-18 | Evans et al. (2014) | bud burst | NA | Eukaryotic protein of unknown function (DUF914) |
| Potri.002G230600 | CO8-18 | Evans et al. (2014) | budset | NA | Eukaryotic protein of unknown function (DUF914) |
| Potri.002G253200 | BB4-top | Evans et al. (2014) | bud burst | NA | CLIP-associated protein |
| Potri.002G255300 | BB4-top | Evans et al. (2014) | budset | NA | TetratriCOpeptide repeat (TPR)-like superfamily protein |
| Potri.002G255400 | BB4-top | Evans et al. (2014) | budset | NA | RPM1 interacting protein 13 |
| Potri.002G255400 | BB4-top | Evans et al. (2014) | height | NA | RPM1 interacting protein 13 |
| Potri.002G256100 | BB4-top | Evans et al. (2014) | height | NA | |
| Potri.002G256200 | BB4-top | Evans et al. (2014) | budset | NA | Exostosin family protein |
| Potri.002G256600 | BB4-top | Evans et al. (2014) | budset | NA | indole-3-acetic acid inducible 11 |
| Potri.002G258300 | BB4-top | Evans et al. (2014) | bud burst | NA | |
| Potri.002G262200 | BB4-top | Evans et al. (2014) | bud burst | NA | |
| Potri.002G263500 | BB4-top | Evans et al. (2014) | bud burst | NA | |

| | | | | | |
|---|---|---|---|---|---|
| Potri.002G263700 | LS5-17, BB4-top | Evans et al. (2014) | budset | NA | |
| Potri.002G263800 | LS5-17 | Evans et al. (2014) | bud burst | NA | cytochrome P450, family 721, subfamily A, polypeptide 1 |
| Potri.002G263900 | LS5-17 | Evans et al. (2014) | bud burst | NA | Plant self-inCOmpatibility protein S1 family |
| Potri.002G264000 | LS5-17 | Evans et al. (2014) | bud burst | NA | |
| Potri.003G044800 | CO3-18 | Evans et al. (2014) | bud burst | NA | |
| Potri.003G044800 | CO3-18 | Evans et al. (2014) | height | NA | |
| Potri.003G075600 | BS2 | Evans et al. (2014) | bud burst | NA | PLATZ transcription factor family protein |
| Potri.003G075600 | BS2 | Evans et al. (2014) | budset | NA | PLATZ transcription factor family protein |
| Potri.003G111700 | LS2-17 | Evans et al. (2014) | bud burst | NA | Phosphatidic acid phosphatase (PAP2) family protein |
| Potri.003G111700 | LS2-17 | Evans et al. (2014) | height | NA | Phosphatidic acid phosphatase (PAP2) family protein |
| Potri.003G123700 | LS5-18 | Evans et al. (2014) | bud burst | NA | UDP-D-gluCOse/UDP-D-galactose 4-epimerase 1 |
| Potri.003G131500 | LS5-18 | Evans et al. (2014) | budset | NA | Nuclear transport factor 2 (NTF2) family protein |
| Potri.003G141600 | LS5-18 | Evans et al. (2014) | budset | NA | Nuclear transport factor 2 (NTF2) family protein |
| Potri.003G141600 | LS5-18 | Evans et al. (2014) | height | NA | Nuclear transport factor 2 (NTF2) family protein |
| Potri.003G214200 | LS2-18 | Mckown et al. (2014) | 100% Yellowing | 2010 | glucan synthase-like 12 |
| Potri.003G214200 | LS2-18 | Mckown et al. (2014) | Branches | 2009 | glucan synthase-like 12 |
| Potri.003G214200 | LS2-18 | Mckown et al. (2014) | Bud set | 2009 | glucan synthase-like 12 |
| Potri.003G214200 | LS2-18 | Mckown et al. (2014) | Bud set | 2010 | glucan synthase-like 12 |
| Potri.003G214200 | LS2-18 | Mckown et al. (2014) | Bud set | 2008 | glucan synthase-like 12 |
| Potri.003G214200 | LS2-18 | Mckown et al. (2014) | Bud set* | 2009 | glucan synthase-like 12 |
| Potri.003G214200 | LS2-18 | Mckown et al. (2014) | Growth period | 2010 | glucan synthase-like 12 |
| Potri.003G214200 | LS2-18 | Mckown et al. (2014) | Height growth cessation | 2009 | glucan synthase-like 12 |
| Potri.003G214200 | LS2-18 | Mckown et al. (2014) | Leaf drop | 2008 | glucan synthase-like 12 |
| Potri.003G214200 | LS2-18 | Mckown et al. (2014) | Leaf drop | 2009 | glucan synthase-like 12 |
| Potri.003G214200 | LS2-18 | Evans et al. (2014) | budset | NA | glucan synthase-like 12 |
| Potri.004G013300 | BB4-top | Evans et al. (2014) | height | NA | PentatriCOpeptide repeat (PPR) superfamily protein |
| Potri.004G013400 | BB4-top | Mckown et al. (2014) | Bud set | 2010 | arogenate dehydratase 1 |
| Potri.004G013400 | BB4-top | Mckown et al. (2014) | Bud set | 2009 | arogenate dehydratase 1 |
| Potri.004G013400 | BB4-top | Mckown et al. (2014) | Bud set* | 2009 | arogenate dehydratase 1 |

| | | | | | |
|---|---|---|---|---|---|
| Potri.004G013400 | BB4-top | Mckown et al. (2014) | Bud set* | 2010 | arogenate dehydratase 1 |
| Potri.004G013400 | BB4-top | Mckown et al. (2014) | Growth period | 2010 | arogenate dehydratase 1 |
| Potri.004G013400 | BB4-top | Mckown et al. (2014) | Height growth cessation | 2009 | arogenate dehydratase 1 |
| Potri.004G013400 | BB4-top | Mckown et al. (2014) | Leaf drop | 2009 | arogenate dehydratase 1 |
| Potri.004G013400 | BB4-top | Mckown et al. (2014) | Leaf drop | 2008 | arogenate dehydratase 1 |
| Potri.004G013400 | BB4-top | Mckown et al. (2014) | Post-bud set period | 2010 | arogenate dehydratase 1 |
| Potri.004G013400 | BB4-top | Evans et al. (2014) | budset | NA | arogenate dehydratase 1 |
| Potri.004G117500 | BB2-top | Evans et al. (2014) | bud burst | NA | PLC-like phosphodiesterases superfamily protein |
| Potri.004G163100 | BB4-top | Evans et al. (2014) | budset | NA | |
| Potri.004G228900 | CO3-17, CO3-18 | Evans et al. (2014) | bud burst | NA | mitotic checkpoint family protein |
| Potri.005G095900 | LS5-17 | Mckown et al. (2014) | Bud set* | 2009 | potassium transporter 1 |
| Potri.005G095900 | LS5-17 | Evans et al. (2014) | bud burst | NA | potassium transporter 1 |
| Potri.005G096000 | LS5-17 | Evans et al. (2014) | bud burst | NA | |
| Potri.005G096100 | LS5-17 | Evans et al. (2014) | bud burst | NA | |
| Potri.005G143400 | BB4-brn, BB4-stb | Evans et al. (2014) | budset | NA | Protein kinase superfamily protein |
| Potri.005G151100 | LS5-17 | Evans et al. (2014) | height | NA | Zinc knuckle (CCHC-type) family protein |
| Potri.005G151800 | BB4-stb | Evans et al. (2014) | budset | NA | Di-gluCOse binding protein with Kinesin motor domain |
| Potri.006G060800 | BB4-stb | Evans et al. (2014) | budset | NA | serine-rich protein-related |
| Potri.006G200200 | LS5-17 | Evans et al. (2014) | bud burst | NA | RING-H2 group F2A |
| Potri.006G200200 | LS5-17 | Evans et al. (2014) | budset | NA | RING-H2 group F2A |
| Potri.006G214400 | LS2-17 | Evans et al. (2014) | bud burst | NA | Pectin lyase-like superfamily protein |
| Potri.007G018300 | LS5-17 | Evans et al. (2014) | budset | NA | |
| Potri.007G018400 | LS5-17 | Evans et al. (2014) | height | NA | Cytochrome P450 superfamily protein |
| Potri.007G045100 | LS5-17 | Evans et al. (2014) | bud burst | NA | Subtilisin-like serine endopeptidase family protein |
| Potri.008G197100 | BB2-top | Evans et al. (2014) | budset | NA | Major facilitator superfamily protein |
| Potri.009G021500 | LS2-17 | Evans et al. (2014) | height | NA | calmodulin 5 |
| Potri.009G050000 | LS5-18 | Evans et al. (2014) | budset | NA | hexokinase-like 1 |
| Potri.009G092400 | LS5-18 | Evans et al. (2014) | budset | NA | Protein of unknown function (DUF620) |
| Potri.009G093500 | BB2-stt | Evans et al. (2014) | budset | NA | 1,2-alpha-L-fuCOsidases |
| Potri.009G095800 | BB2-stt | Evans et al. (2014) | height | NA | GroES-like zinc-binding alCOhol dehydrogenase family protein |
| Potri.009G162800 | CO8-17 | Evans et al. (2014) | bud burst | NA | Protein of unknown function (DUF3049) |

| | | | | | |
|---|---|---|---|---|---|
| Potri.010G016300 | BB4-stb | Evans et al. (2014) | bud burst | NA | Raffinose synthase family protein |
| Potri.010G016400 | BB4-stb | Evans et al. (2014) | height | NA | Raffinose synthase family protein |
| Potri.010G066900 | BB4-stb | Evans et al. (2014) | bud burst | NA | proton gradient regulation 5 |
| Potri.010G167200 | CO3-17 | Evans et al. (2014) | budset | NA | expansin A1 |
| Potri.010G180000 | CO3-17 | Evans et al. (2014) | budset | NA | HXXXD-type acyl-transferase family protein |
| Potri.010G180100 | CO3-17 | Evans et al. (2014) | budset | NA | |
| Potri.010G180100 | CO3-17 | Evans et al. (2014) | height | NA | |
| Potri.010G180300 | CO3-17 | Evans et al. (2014) | budset | NA | Cysteine proteinases superfamily protein |
| Potri.010G180400 | CO3-17 | Evans et al. (2014) | budset | NA | 2-phosphoglyCOlate phosphatase 1 |
| Potri.010G180500 | CO3-17 | Evans et al. (2014) | budset | NA | UDP-galactose transporter 6 |
| Potri.010G180800 | LS5-17 | Evans et al. (2014) | budset | NA | Calcineurin-like metallo-phosphoesterase superfamily protein |
| Potri.010G181000 | LS5-17 | Evans et al. (2014) | budset | NA | Integrase-type DNA-binding superfamily protein |
| Potri.010G181000 | LS5-17 | Evans et al. (2014) | height | NA | Integrase-type DNA-binding superfamily protein |
| Potri.012G070600 | CO3-17 | Evans et al. (2014) | bud burst | NA | Protein of unknown function (DUF1118) |
| Potri.012G070700 | CO3-17 | Evans et al. (2014) | bud burst | NA | nitrate transporter 1:2 |
| Potri.012G077400 | LS5-17 | Evans et al. (2014) | budset | NA | Xanthine/uracil permease family protein |
| Potri.012G105000 | LS5-17 | Evans et al. (2014) | bud burst | NA | Polynucleotide adenylyltransferase family protein |
| Potri.012G105000 | LS5-17 | Evans et al. (2014) | budset | NA | Polynucleotide adenylyltransferase family protein |
| Potri.012G105200 | LS5-17 | Evans et al. (2014) | bud burst | NA | AAA-type ATPase family protein |
| Potri.013G001300 | BS7 | Evans et al. (2014) | budset | NA | phytochrome interacting factor 3 |
| Potri.013G003100 | BS7 | Evans et al. (2014) | height | NA | Putative endonuclease or glyCOsyl hydrolase |
| Potri.013G003200 | BS7 | Evans et al. (2014) | budset | NA | Putative endonuclease or glyCOsyl hydrolase |
| Potri.013G026100 | CO8-17, LS5-17 | Evans et al. (2014) | bud burst | NA | NAD(P)-binding Rossmann-fold superfamily protein |
| Potri.013G026100 | CO8-17, LS5-17 | Evans et al. (2014) | budset | NA | NAD(P)-binding Rossmann-fold superfamily protein |
| Potri.013G026100 | CO8-17, LS5-17 | Evans et al. (2014) | height | NA | NAD(P)-binding Rossmann-fold superfamily protein |
| Potri.013G090400 | LS5-17 | Evans et al. (2014) | bud burst | NA | WRKY DNA-binding protein 55 |
| Potri.013G090500 | LS5-17 | Evans et al. (2014) | bud burst | NA | Protein kinase family protein with ARM repeat domain |
| Potri.013G090800 | LS5-17 | Evans et al. (2014) | height | NA | SNF1 kinase homolog 10 |

| | | | | | |
|---|---|---|---|---|---|
| Potri.013G094000 | BB2-18 | Evans et al. (2014) | budset | NA | S-locus lectin protein kinase family protein |
| Potri.013G094000 | BB2-18 | Evans et al. (2014) | height | NA | S-locus lectin protein kinase family protein |
| Potri.013G100600 | LS2-17 | Evans et al. (2014) | bud burst | NA | Nuclear transport factor 2 (NTF2) family protein |
| Potri.013G100600 | LS2-17 | Evans et al. (2014) | height | NA | Nuclear transport factor 2 (NTF2) family protein |
| Potri.013G118700 | CO3-17 | Evans et al. (2014) | bud burst | NA | UDP-gluCOsyl transferase 78D2 |
| Potri.013G118800 | CO3-17 | Evans et al. (2014) | bud burst | NA | |
| Potri.013G118900 | CO3-17 | Evans et al. (2014) | bud burst | NA | global transcription factor group E4 |
| Potri.013G130600 | BB4-top | Evans et al. (2014) | bud burst | NA | PentatriCOpeptide repeat (PPR) superfamily protein |
| Potri.013G130700 | BB4-top | Evans et al. (2014) | bud burst | NA | |
| Potri.013G147800 | CO3-17 | Evans et al. (2014) | bud burst | NA | PapD-like superfamily protein |
| Potri.013G148000 | CO3-17 | Evans et al. (2014) | height | NA | Wound-responsive family protein |
| Potri.014G000100 | BB4-stb | Evans et al. (2014) | height | NA | zinc ion binding;nucleic acid binding |
| Potri.015G049100 | LS5-17 | Mckown et al. (2014) | Leaf drop | 2009 | glucuronidase 2 |
| Potri.015G049100 | LS5-17 | Evans et al. (2014) | budset | NA | glucuronidase 2 |
| Potri.015G050300 | LS5-17 | Evans et al. (2014) | budset | NA | Transmembrane Fragile-X-F-associated protein |
| Potri.015G099200 | BB4-18 | Evans et al. (2014) | bud burst | NA | WRKY DNA-binding protein 75 |
| Potri.015G099400 | BB4-18 | Evans et al. (2014) | bud burst | NA | |
| Potri.016G007000 | BS7 | Evans et al. (2014) | height | NA | |
| Potri.016G030000 | BB4-18 | Evans et al. (2014) | bud burst | NA | Leucine-rich repeat protein kinase family protein |
| Potri.016G030100 | BB4-18 | Evans et al. (2014) | bud burst | NA | syntaxin of plants 51 |
| Potri.016G030200 | BB4-18 | Evans et al. (2014) | bud burst | NA | syntaxin of plants 52 |
| Potri.016G030400 | BB4-18 | Evans et al. (2014) | bud burst | NA | Transducin/WD40 repeat-like superfamily protein |
| Potri.016G030500 | BB4-17, BB4-18 | Evans et al. (2014) | bud burst | NA | DNA/RNA helicase protein |
| Potri.016G030600 | BB4-17, BB4-18 | Evans et al. (2014) | bud burst | NA | DNA/RNA helicase protein |
| Potri.016G033300 | LS2-17 | Evans et al. (2014) | bud burst | NA | Transcription elongation factor (TFIIS) family protein |
| Potri.016G033300 | LS2-17 | Evans et al. (2014) | budset | NA | Transcription elongation factor (TFIIS) family protein |
| Potri.016G033400 | LS2-17, LS2-18 | Evans et al. (2014) | budset | NA | |
| Potri.016G033500 | LS2-17, LS2-18 | Evans et al. (2014) | budset | NA | DNA/RNA helicase protein |
| Potri.016G033600 | LS2-17, LS2-18 | Evans et al. (2014) | budset | NA | DNA/RNA helicase protein |
| Potri.016G043700 | CO3-18 | Evans et al. (2014) | bud burst | NA | alpha carbonic anhydrase 4 |
| Potri.016G043800 | CO3-18 | Evans et al. (2014) | height | NA | eukaryotic translation initiation factor 3E |

| | | | | | |
|---|---|---|---|---|---|
| Potri.016G098100 | LS5-17 | Evans et al. (2014) | bud burst | NA | C2H2-type zinc finger family protein |
| Potri.016G098100 | LS5-17 | Evans et al. (2014) | height | NA | C2H2-type zinc finger family protein |
| Potri.016G137500 | BB4-17 | Evans et al. (2014) | bud burst | NA | VACUOLAR SORTING RECEPTOR 2 |
| Potri.016G137500 | BB4-17 | Evans et al. (2014) | height | NA | VACUOLAR SORTING RECEPTOR 2 |
| Potri.016G137700 | BB4-17 | Evans et al. (2014) | budset | NA | Lung seven transmembrane receptor family protein |
| Potri.016G137800 | BB4-17 | Evans et al. (2014) | budset | NA | Protein of unknown function, DUF599 |
| Potri.017G023300 | BS2 | Evans et al. (2014) | budset | NA | Protein of unknown function (DUF3755) |
| Potri.017G039700 | BS2 | Evans et al. (2014) | height | NA | cysteine-rich RLK (RECEPTOR-like protein kinase) 16 |
| Potri.017G092200 | CO8-17 | Evans et al. (2014) | bud burst | NA | Ribosomal protein S19e family protein |
| Potri.017G093600 | CO8-17 | Evans et al. (2014) | height | NA | RS2-interacting KH protein |
| Potri.017G148900 | BB2-18 | Evans et al. (2014) | budset | NA | Tic22-like family protein |
| Potri.018G008200 | BB4-brn | Evans et al. (2014) | bud burst | NA | |
| Potri.018G082100 | BB4-stt | Evans et al. (2014) | budset | NA | Protein of unknown function (DUF502) |
| Potri.018G082300 | BB4-stt | Evans et al. (2014) | bud burst | NA | |
| Potri.018G082300 | BB4-stt | Evans et al. (2014) | budset | NA | |
| Potri.019G063400 | LS5-17 | Evans et al. (2014) | bud burst | NA | FAD-binding Berberine family protein |

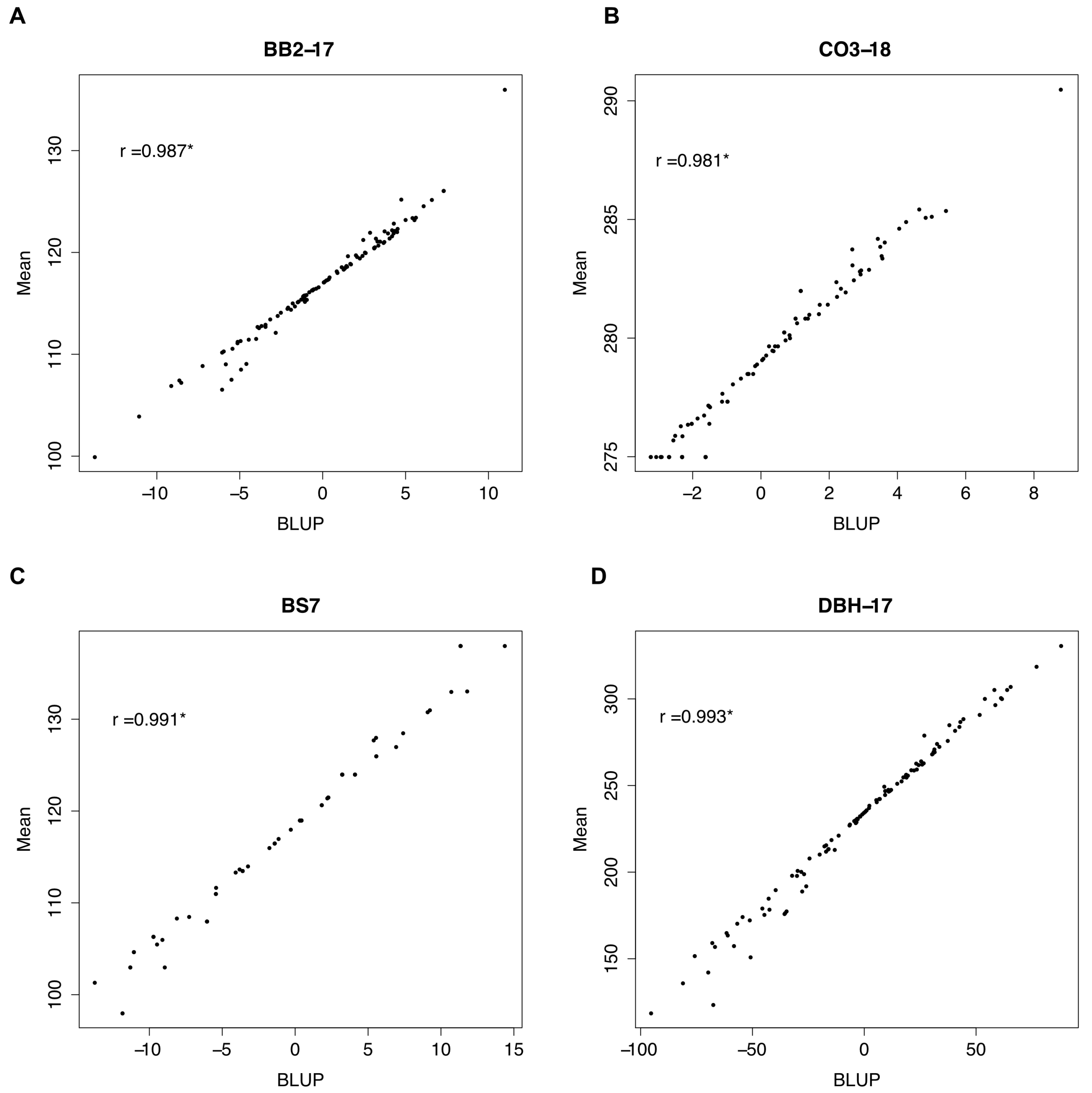


Figure S1: Pearson’s correlation between BLUP and mean values of A) bud burst initation in year 2017 (BB2-17), B) autumn coloring initiation in year 2018 (CO3-18), C) bud set completion (BS7) and D) diameter at breast height (DBH-17). * $p < 2.2^{-16}$.

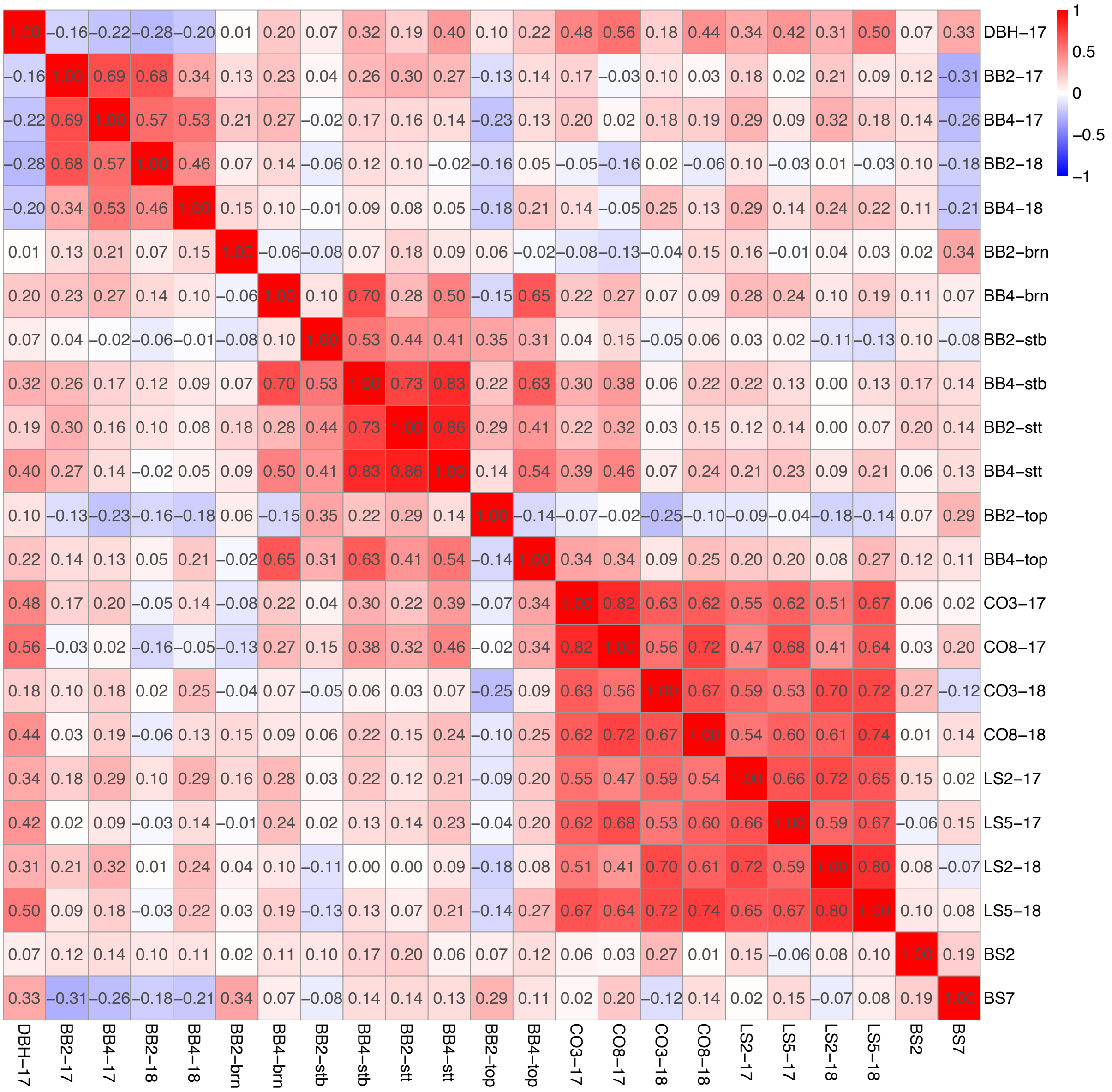


Figure S2: Spearman's correlation matrix and heatmap of all our chosen 23 traits.

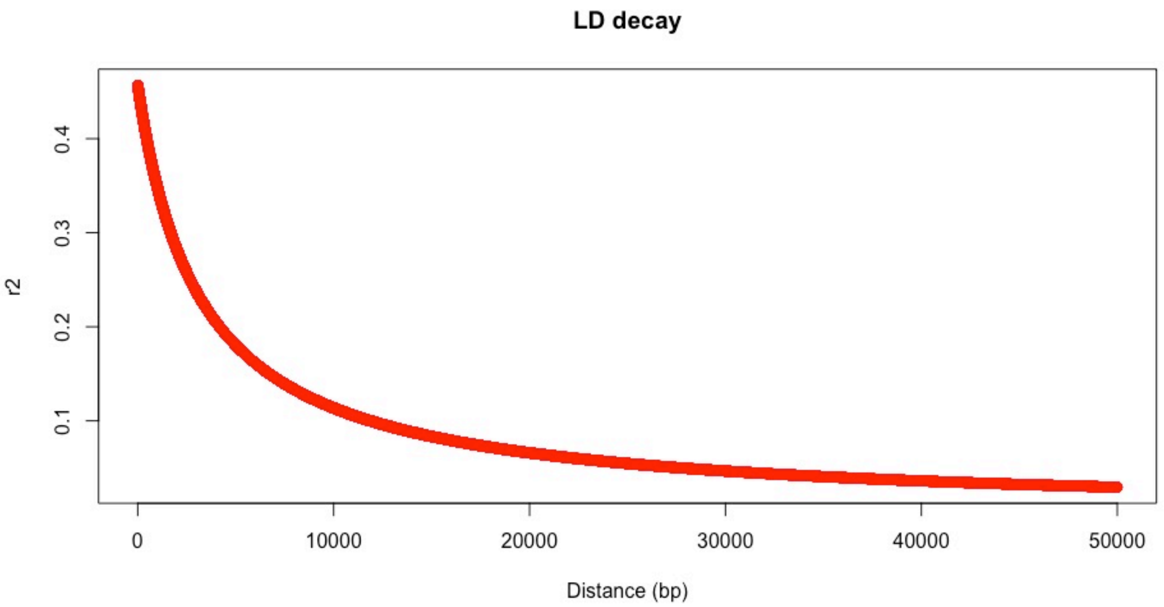


Figure S3: Level of LD as function of distance.

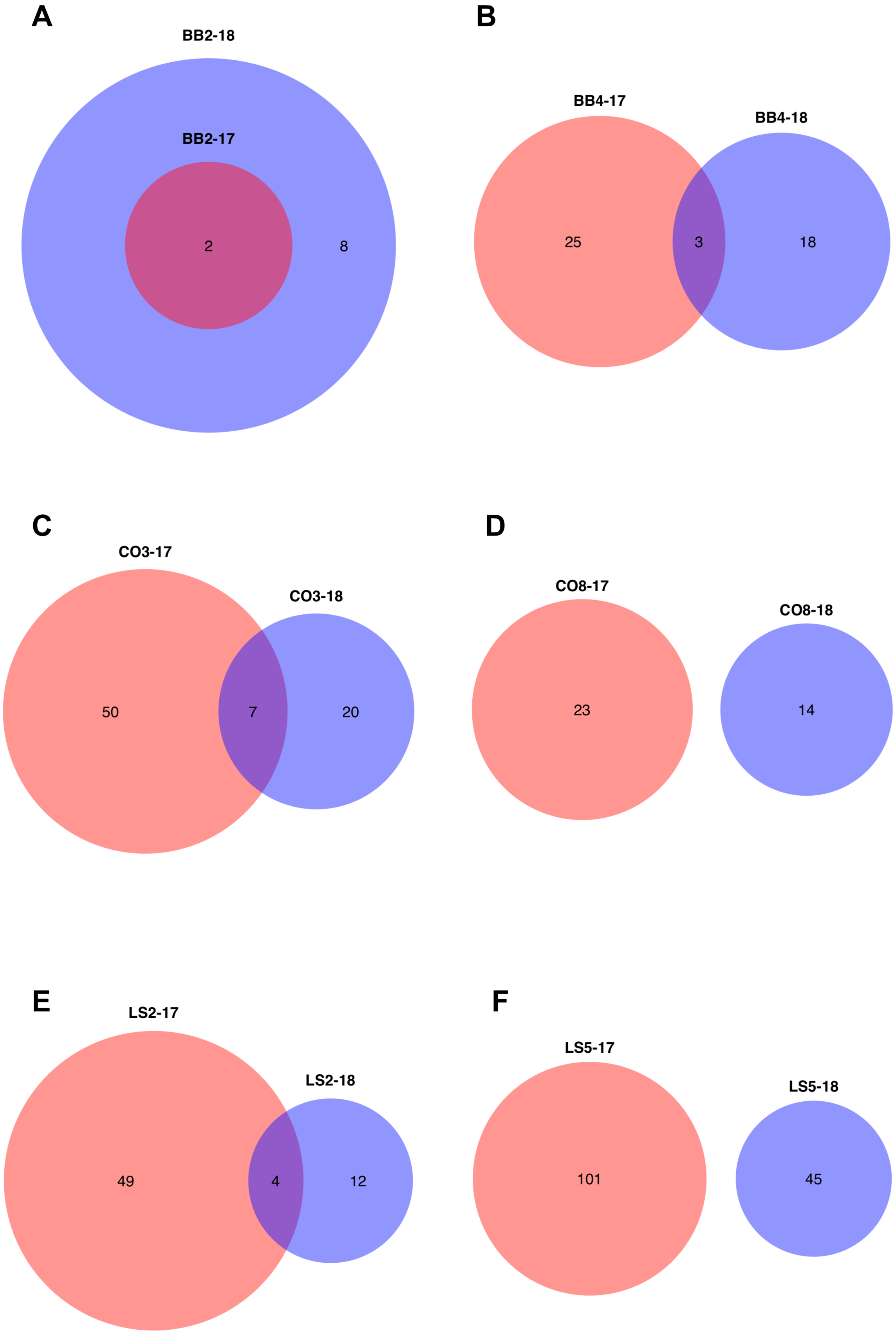
A
BB2-18
BB2-17
2
8
B
BB4-17
BB4-18
25
3
18
C
CO3-17
CO3-18
50
7
20
D
CO8-17
CO8-18
23
14
E
LS2-17
LS2-18
49
4
12
F
LS5-17
LS5-18
101
45

Figure S4: Venn diagram of candidate genes for both year for A) bud burst initiation (BB2), B) bud burs completion (BB4) C) autumn coloring initiation (CO3), D) autumn coloring completion (CO8) E) leaf shed initiation (LS2) and F) leaf shed completion (LS5).
